# Impaired local intrinsic immunity to SARS-CoV-2 infection in severe COVID-19

**DOI:** 10.1101/2021.02.20.431155

**Authors:** Carly G. K. Ziegler, Vincent N. Miao, Anna H. Owings, Andrew W. Navia, Ying Tang, Joshua D. Bromley, Peter Lotfy, Meredith Sloan, Hannah Laird, Haley B. Williams, Micayla George, Riley S. Drake, Taylor Christian, Adam Parker, Campbell B. Sindel, Molly W. Burger, Yilianys Pride, Mohammad Hasan, George E. Abraham, Michal Senitko, Tanya O. Robinson, Alex K. Shalek, Sarah C. Glover, Bruce H. Horwitz, Jose Ordovas-Montanes

**Author notes:** these authors contributed equally (Carly G. K. Ziegler, Vincent N. Miao, Anna H. Owings, Andrew W. Navia, Ying Tang, Joshua D. Bromley). these senior authors contributed equally (Alex K. Shalek, Sarah C. Glover, Bruce H. Horwitz, Jose Ordovas-Montanes). Correspondence to: Jose Ordovas-Montanes, Bruce Horwitz, Sarah C. Glover, and Alex K. Shalek.

## Abstract

Infection with SARS-CoV-2, the virus that causes COVID-19, can lead to severe lower respiratory illness including pneumonia and acute respiratory distress syndrome, which can result in profound morbidity and mortality. However, many infected individuals are either asymptomatic or have isolated upper respiratory symptoms, which suggests that the upper airways represent the initial site of viral infection, and that some individuals are able to largely constrain viral pathology to the nasal and oropharyngeal tissues. Which cell types in the human nasopharynx are the primary targets of SARS-CoV-2 infection, and how infection influences the cellular organization of the respiratory epithelium remains incompletely understood. Here, we present nasopharyngeal samples from a cohort of 35 individuals with COVID-19, representing a wide spectrum of disease states from ambulatory to critically ill, as well as 23 healthy and intubated patients without COVID-19. Using standard nasopharyngeal swabs, we collected viable cells and performed single-cell RNA-sequencing (scRNA-seq), simultaneously profiling both host and viral RNA. We find that following infection with SARS-CoV-2, the upper respiratory epithelium undergoes massive reorganization: secretory cells diversify and expand, and mature epithelial cells are preferentially lost. Further, we observe evidence for deuterosomal cell and immature ciliated cell expansion, potentially representing active repopulation of lost ciliated cells through coupled secretory cell differentiation. Epithelial cells from participants with mild/moderate COVID-19 show extensive induction of genes associated with anti-viral and type I interferon responses. In contrast, cells from participants with severe lower respiratory symptoms appear globally muted in their anti-viral capacity, despite substantially higher local inflammatory myeloid populations and equivalent nasal viral loads. This suggests an essential role for intrinsic, local epithelial immunity in curbing and constraining viral-induced pathology. Using a custom computational pipeline, we characterized cell-associated SARS-CoV-2 RNA and identified rare cells with RNA intermediates strongly suggestive of active replication. Both within and across individuals, we find remarkable diversity and heterogeneity among SARS-CoV-2 RNA+ host cells, including developing/immature and interferon-responsive ciliated cells, *KRT13+* “hillock”-like cells, and unique subsets of secretory, goblet, and squamous cells. Finally, SARS-CoV-2 RNA+ cells, as compared to uninfected bystanders, are enriched for genes involved in susceptibility (e.g., *CTSL*, *TMPRSS2*) or response (e.g., *MX1*, *IFITM3*, *EIF2AK2*) to infection. Together, this work defines both protective and detrimental host responses to SARS-CoV-2, determines the direct viral targets of infection, and suggests that failed anti-viral epithelial immunity in the nasal mucosa may underlie the progression to severe COVID-19.

## INTRODUCTION

The novel coronavirus clade SARS-CoV-2 emerged in late 2019 and has quickly led to one of the most devastating global pandemics in modern history. Similar to other successful respiratory viruses, high replication within the nasopharynx^1,2^ and viral shedding by asymptomatic or presymptomatic individuals contributes to high transmissibility^3,4^ and rapid community spread^5–7^. COVID-19, the disease caused by SARS-CoV-2 infection, occurs in a fraction of those infected by the virus and carries profound morbidity and mortality. The clinical pictures of COVID-19 vary widely – from some individuals who experience few mild symptoms to some with prolonged and severe disease characterized by pneumonia, acute respiratory distress syndrome, and diverse systemic effects impacting various other tissues^8,9^. To facilitate effective preventative and therapeutic strategies for COVID-19, differentiating the host protective mechanisms that support rapid viral clearance and limit disease severity from those that drive severe and fatal outcomes is essential.

Rapid mobilization of the scientific community and a commitment to open data sharing early in the COVID-19 pandemic enabled researchers across the globe to study SARS-CoV-2 and build initial models of disease pathogenesis^10–12^. By analogy to related human betacoronaviruses^13^, we currently understand viral tropism and disease progression to begin with SARS-CoV-2 entry through the mouth or nares where it initially replicates within epithelial cells of the human nasopharynx, generating an upper respiratory infection over several days^14^. A subset of patients develop symptoms of lower respiratory, where a combination of inflammatory immune responses and direct viral-mediated pathogenesis can lead to diffuse damage to distal airways, alveoli, and vasculature^15,16^. Recent studies have mapped the host immune profiles associated with different stages along the COVID-19 disease trajectory. Reproducible immune correlates of severe COVID-19 include prolonged detection of proinflammatory cytokines such as IL-6, TNFα, and IL-8, diminished type I and type III interferon, and marked lymphopenia, as well as mixed evidence for immune exhaustion and dysfunctional myeloid populations^17–25^. These reports have measured host inflammatory and immune signatures in peripheral blood, which may only partially reflect the immune status within virally targeted tissues^26,27^. To date, few studies have directly addressed the impact of SARS-CoV-2 infection on the respiratory epithelium of the human upper airways, or examined how this may relate to aberrant inflammatory or anti-viral signaling described in the periphery.

A question central to understanding SARS-CoV-2-induced disease pathology is the precise identity of the direct cellular targets of viral infection within human respiratory tissues. Early in the pandemic, multiple groups conducted meta-analyses of existing single-cell RNA-sequencing (scRNA-seq) datasets from diverse host tissues to map potential SARS-CoV-2 tropism based on *ACE2* expression and co-expression of host proteases required for spike protein cleavage^28–32^. Together, these studies nominated putative SARS-CoV-2-targeted cells within the oropharyngeal, nasal, and upper airway tissues including subsets of ciliated, secretory, and goblet cells, and within the lung parenchyma, type II pneumocytes. Indeed, a study jointly collecting nasopharyngeal and bronchoalveolar lavage samples from a cohort of COVID-19 patients identified rare SARS-CoV-2 RNA-containing cells assigned to ciliated and secretory cell types^33^. Further work using human tissues at autopsy found infected ciliated cells lining the trachea and distal airways within the lungs^34–36^. In vitro studies have illustrated the capacity of SARS-CoV-2 to infect myriad organoid and air-liquid interface models of tissues providing important lessons about mechanisms of entry and the anti-viral or inflammatory responses induced^37–44^. However, the precise early targets for SARS-CoV-2 in the nasopharynx, the scope of potential host cells, and the variance in viral tropism across patients and disease courses have yet to be defined. A clearer understanding of viral tropism, how the airway epithelium responds to infection, and the relationship to disease outcome may critically inform future therapeutic or prophylactic strategies.

Further, we currently lack a clear understanding of the host factors responsible for susceptibility versus resistance to viral infection. Researchers have employed *in vitro* systems to assess induction of anti-viral defenses following SARS-CoV-2 infection. Compared to other common respiratory viruses, SARS-CoV-2 appears to elicit poor type I interferon responses in cultured human epithelial cells, and instead skews towards proinflammatory cytokine profiles, in line with observations from human peripheral studies^21,37,38^. To directly assay virally-targeted cell types or tissues *in vivo,* researchers have relied on emerging animal models, including non-human primates^45–47^, hamsters^48,49^, mice^50–53^, and ferrets^54,55^. These animal models vary widely in the severity of SARS-CoV-2-driven disease and associated immunopathology, and incompletely reflect the diversity of viral infection outcomes and natural immune responses within the human population^56^. Recent work leveraging human cohorts has identified enrichment of both inborn errors of type I interferon signaling and the presence of auto-antibodies against type I interferons among patients with severe COVID-19, providing potential explanations for failed or insufficient anti-viral immunity within a subset of severe cases, and further supporting the need for human cohort studies that represent the breadth of host-viral interactions^57–59^.

Here, we present a comprehensive analysis of the cellular phenotypes in the nasal mucosa during early SARS-CoV-2 infection. To achieve this, we developed tissue handling protocols that enabled high-quality scRNA-seq from frozen nasopharyngeal swabs collected from a large patient cohort (n = 58), and created a detailed map of epithelial and immune cell diversity. We found that SARS-CoV-2 infection leads to a dramatic loss of mature ciliated cells, which is associated with secretory cell expansion, differentiation, and the accumulation of deuterosomal cell intermediates – potentially involved in the compensatory repopulation of damaged ciliated epithelium. While we observe broad induction of interferon-responsive and anti-viral genes in cells from individuals with mild/moderate COVID-19, severe COVID-19 is characterized by a dramatically blunted interferon response, and mucosal recruitment of highly inflammatory myeloid populations, which represent the primary sources of tissue pro-inflammatory cytokines including *TNF*, *IL1B*, and *CXCL8*. Further, using unbiased whole-transcriptomic amplification, we map not only host cellular RNA, but also cell-associated SARS-CoV-2 RNA, allowing us to trace viral tropism to specific epithelial subsets and identify host pathways linked with susceptibility or resistance to viral infection. Together, our data suggest that an early intrinsic failure of anti-viral immunity among nasal epithelial cells responding to SARS-CoV-2 infection may predict progression to severe COVID-19.

## RESULTS

### Defining Cellular Diversity in the Human Nasopharyngeal Mucosa

Nasopharyngeal swabs were collected from 58 individuals from the University of Mississippi Medical Center between April and September 2020. This cohort consisted of 35 individuals who had a positive SARS-CoV-2 PCR nasopharyngeal (NP) swab on the day of hospital presentation. A Control group consisted of 15 individuals who were asymptomatic and had a negative SARS-CoV-2 NP PCR, 6 intubated individuals in the intensive care unit without a recent history of COVID-19 and negative SARS-CoV-2 NP PCR, and 2 additional individuals with recent history of COVID-19 and negative SARS-CoV-2 NP PCR, classified as “Convalescent” (**Table 1**, see **Methods** for full inclusion and exclusion criteria). For the purposes of this study a second NP swab was collected within 3 days of presentation. Using the World Health Organization (WHO) guidelines for stratification and classification of COVID-19 severity based on the level of maximum required respiratory support^60^: 14 of the individuals were considered COVID-19 mild/moderate (WHO score 1-5) and 21 had severe COVID-19 (WHO score 6-8, see **Methods, Table 1, Supplementary Figures 1A, 1B**). Nasopharyngeal samples were obtained by a trained healthcare provider and rapidly cryopreserved to maintain cellular viability (**Figure 1A, Supplementary Figure 1C**). Swabs were later processed to recover single-cell suspensions (mean ± SEM: 57,000 ± 15,000 total cells recovered per swab), before generating single-cell transcriptomes using Seq-Well S^3 61,62^.

**Figure 1.**
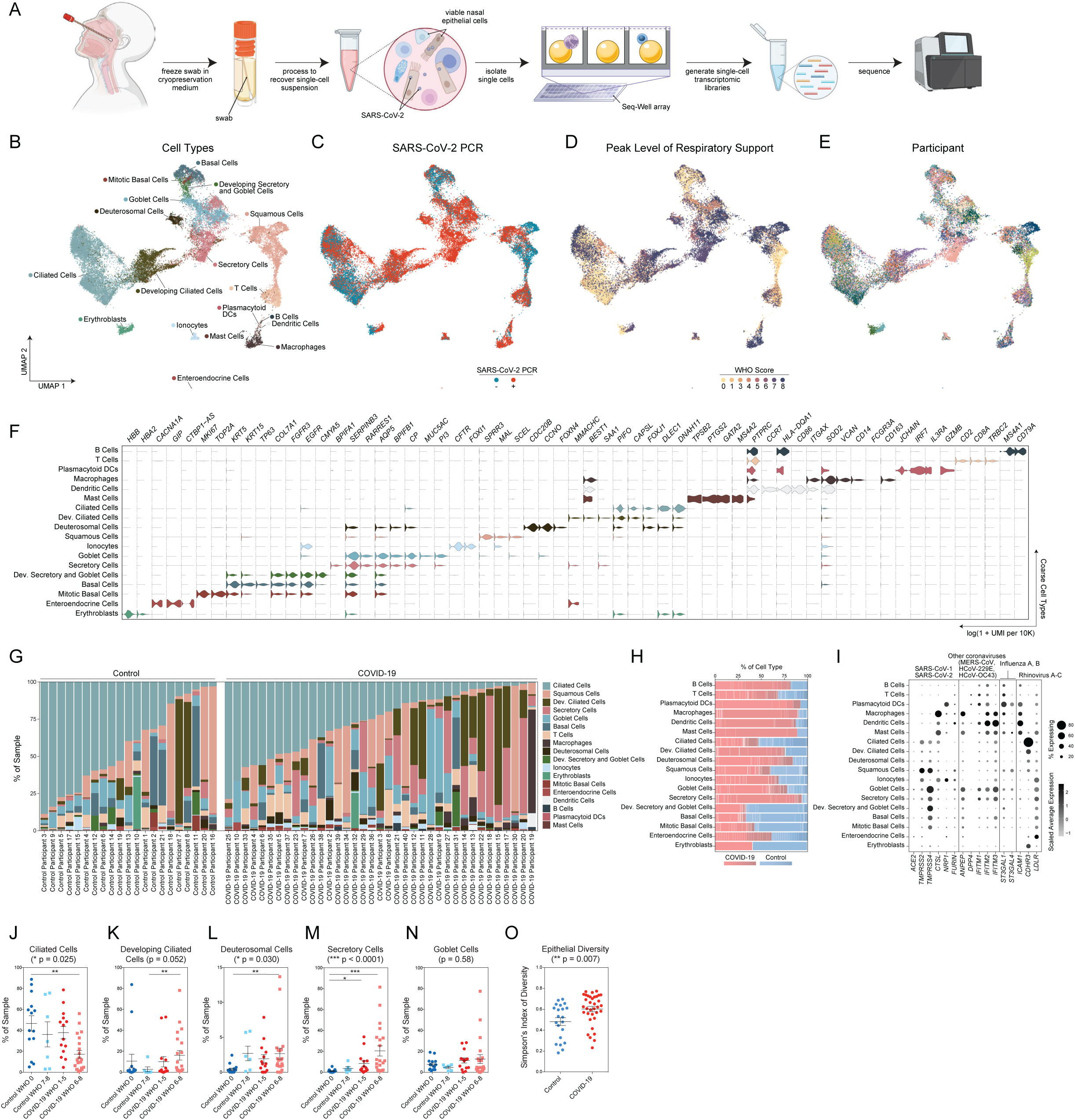
Cellular Composition of Human Nasopharyngeal Mucosa. **A.** Schematic of method for viable cryopreservation of nasopharyngeal swabs, cellular isolation, and scRNA-seq using Seq-Well S^3^ (created with BioRender). **B.** UMAP of 32,588 single-cell transcriptomes from all participants, colored by cell type (following iterative Louvain clustering). **C.** UMAP as in **B**, colored by SARS-CoV-2 diagnostic PCR status. **D.** UMAP as in **B**, colored by peak level of respiratory support (WHO COVID-19 severity scale). **E.** UMAP as in **B**, colored by participant. **F.** Violin plots (log(1+normalized UMI per 10k)) of cluster marker genes (FDR < 0.01) for coarse cell type annotations (as in **B**). **G.** Proportional abundance of coarse cell types by participant (ordered within each group by increasing ciliated cell abundance). **H.** Proportional abundance of participants by coarse cell types. Shades of red: COVID-19. Shades of blue: Control. **I.** Expression of entry factors for SARS-CoV-2 and other common upper respiratory viruses. Dot size represents fraction of cell type (rows) expressing a given gene (columns). Dot hue represents scaled average expression by gene column. **J.** Proportion of ciliated cells by sample. Statistical test above graph represents Kruskal-Wallis test results across all groups (following FDR correction across cell types). Statistical significance asterisks within box represent results from Dunn’s post-hoc testing. * p < 0.05, ** p < 0.01, *** p < 0.001. **K.** Proportion of developing ciliated cells by sample. **L.** Proportion of deuterosomal cells by sample. **M.** Proportion of secretory cells by sample. **N.** Proportion of goblet cells by sample. **O.** Simpson’s Diversity index (plotted as 1-D, where increasing values represent higher diversity) across epithelial cell types in COVID-19 vs. Control. Significance by student’s t-test. Lines represent mean and S.E.M. *See also Figure S1, Table S1*

**Table 1.** Participant characteristics.

|  | Control<br>(WHO score 0) | Intubated Control<br>(WHO score 7-8) | COVID-19 m/m<br>(WHO score 1-5) | COVID-19 severe<br>(WHO score 6-8) | COVID-19 conv.<br>(WHO score 0) |
| --- | --- | --- | --- | --- | --- |
| Case number | 25.9% (15/58) | 10.3% (6/58) | 24.1% (14/58) | 36.2 (21/58) | 3.4% (2/58) |
| Age (years) |  |  |  |  |  |
| Minimum | 27 | 33 | 19 | 28 | 20 |
| Median (IQR) | 58 (16) | 65.5 (31) | 49.5 (17.8) | 62 (13) | N/A |
| Maximum | 73 | 71 | 69 | 84 | 57 |
| Sex |  |  |  |  |  |
| Female | 60% (9/15) | 16.7% (1/6) | 42.9% (6/14) | 47.6% (10/21) | 50% (1/2) |
| Male | 40% (6/15) | 83.3% (5/6) | 57.1% (8/14) | 52.4% (11/21) | 50% (1/2) |
| Ethnicity |  |  |  |  |  |
| Hispanic | 0% (0/15) | 0% (0/6) | 0% (0/14) | 4.8% (1/21) | 0% (0/2) |
| Not Hispanic | 100% (15/15) | 100% (6/6) | 100% (14/14) | 95.2% (20/21) | 100% (2/2) |
| Race |  |  |  |  |  |
| Black/African American | 66.7% (10/15) | 66.7% (4/6) | 71.4% (10/14) | 61.9% (13/21) | 50% (1/2) |
| White | 33.3% (5/15) | 33.3% (2/6) | 28.6% (4/14) | 23.8% (5/21) | 50% (1/2) |
| American Indian | 0% (0/15) | 0% (0/6) | 0% (0/14) | 14.3% (3/21) | 0% (0/2) |
| BMI |  |  |  |  |  |
| Median (IQR) | 37.5 (14.4) | 30.5 (18.1) | 23.0 (11.6) | 31.9 (14.2) | 40.7 |
| Pre-existing conditions |  |  |  |  |  |
| Diabetes | 40% (6/15) | 33.3% (2/6) | 28.6% (4/14) | 71.4% (15/21) | 0% (0/2) |
| Chronic kidney disease | 6.7% (1/15) | 0% (0/6) | 7.1% (1/14) | 19.0% (4/21) | 0% (0/2) |
| Congestive heart failure | 6.7% (1/15) | 16.7% (1/6) | 0% (0/14) | 4.8% (1/21) | 0% (0/2) |
| Lung disorder | 6.7% (1/15) | 16.7% (1/6) | 28.6% (4/14) | 38.1% (8/21) | 0% (0/2) |
| Hypertension | 86.7% (13/15) | 50% (3/6) | 42.9% (6/14) | 81.0% (17/21) | 0% (0/2) |
| IBD | 13.3% (2/15) | 0% (0/6) | 0% (0/14) | 0% (0/21) | 50% (1/2) |
| Treatment |  |  |  |  |  |
| Corticosteroids | N/A | 33.3% (2/6) | 42.9% (6/14) | 66.7% (14/21) | N/A |
| Remdesivir | N/A | 0% (0/6) | 42.9% (6/14) | 85.7% (18/21) | N/A |
| 28-day mortality | 0% (0/15) | 33.3% (2/6) | 0% (0/14) | 76.2% (16/21) | 0% (0/2) |

m/m: mild/moderate
conv: convalescent
IQR: inter-quartile range
BMI: body mass index
IBD: inflammatory bowel disease

Among all COVID-19 and Control samples, we recovered 32,871 genes across 32,588 cells (following filtering and quality control), with an average recovery of 562 ± 69 cells per swab (mean ± SEM). We found roughly equivalent transcriptomic quality following uniform preprocessing steps between COVID-19 and Control participants, despite variability in cellular recovery between participants (**Supplementary Figures 1D, 1E**). Following dimensionality reduction and clustering approaches to resolve individual cell types and cell states, we annotated 18 clusters corresponding to distinct cell types across immune and epithelial identities (**Figures 1B-E, Supplementary Table 1)**. Consistent with the use of nasal swabs for cell collection, we did not recover stromal cell populations such as endothelial cells, fibroblasts, or pericytes, which were found in previous scRNA-seq datasets from nasal epithelial surgical samples^63–65^. Among epithelial cell types, we readily identified basal cells by their expression of canonical marker genes including *TP63, KRT15,* and *KRT5*, as well as mitotic basal cells based on the added expression of genes involved in the cell cycle such as *MKI67* and *TOP2A* (**Figure 1F**). We resolved large populations of both secretory cells and goblet cells, identified by expression of *KRT7*, *CXCL17, F3, AQP5,* and *CP*. Despite strong transcriptional similarity between secretory and goblet cells, we distinguished between them based on expression of *MUC5AC*, which defines goblet cells, and *BPIFA1*, which we found primarily expressed within secretory cell types and diminished in *MUC5AC* high cells. We also designated a small population of cells “developing secretory and goblet cells” based on their lower expression of classic secretory/goblet cell genes, as well as persistent expression of some basal cell markers (e.g., persistent *COL7A1* and *DST* expression, but diminishing *KRT5*, *KRT15* expression). We also resolved a population of ionocytes, a recently-identified specialized subtype of secretory cell involved in regulating mucus viscosity within respiratory epithelia, defined by expression of *FOXI1, FOXI2*, and *CFTR*^66,67^. Squamous cells were identified by their expression of *SCEL*, as well as multiple SPRR genes, and potentially derive from the squamous epithelium of the anterior nose or posterior pharynx. We also recovered a very small population of cells we term “enteroendocrine cells”, based on unique expression of gastric inhibitory polypeptide (*GIP*), which is typically produced by intestinal and gastric enteroendocrine cells and *LGR5*, which classically marks stem cell populations in the gastrointestinal mucosa^68^.

Ciliated cells, defined by expression of transcription factor *FOXJ1* as well as numerous genes involved in the formation of cilia, e.g., *DLEC1, DNAH11,* and *CFAP43*, were the most numerous epithelial cell type recovered in this dataset. Similar to intermediate/developing cells of the secretory and goblet lineage, we also identified two populations of precursor ciliated cells. One, termed “developing ciliated cells”, which expressed canonical ciliated cell genes such as *FOXJ1*, *CAPSL*, and *PIFO* at lower levels than mature ciliated cells and lacked expression of cilia-forming genes. We also identified a cluster defined by expression of *DEUP1*, which is critical for centriole amplification as a precursor to cilium assembly, as well as *CCNO, CDC20B, FOXN4,* and *HES6*. This profile matches that of a recently-defined cell type termed deuterosomal cells, which represent a ciliated cell precursor cell type arising from secretory cell/goblet cell differentiation^64^.

Immune cells were a minority of recovered cells, yet we resolved multiple distinct clusters and cell types, representing major myeloid and lymphoid populations. Among lymphoid cells, we recovered T cells, identified by *CD3E, CD2,* and *TRBC2* expression, and B cells, identified by *MS4A1, CD79A,* and *CD79B* expression. Among myeloid cell types, we recovered a large population of macrophages (*CD14, FCGR3A, VCAN*), dendritic cells (*CCR7, CD86*), and plasmacytoid DCs (*IRF7, IL3RA*). Relative to true tissue-resident abundances, we under-recovered granulocyte populations, likely due to the intrinsic fragility of these cell types and the cryopreservation methods required in our sample pipeline (**Supplementary Figure 1F**). We recovered a very small population of mast cells, defined by expression of *GATA2, TPSB2,* and *PTGS2.* Each cell type is represented by cells from numerous participants, and from each participant we recovered a diversity of cell types and states, though the cellular composition is highly variable between distinct individuals (**Figure 1G, 1H**).

We directly tested whether cell types collected from nasal swabs following cryopreservation were representative of cellular composition extracted from a freshly swabbed nasal epithelium, or if certain cell types were lost during freezing (**Supplementary Figure 1G-1L**). Recovery of viable cells, technical metrics of single-cell library quality, and cellular proportions after clustering and analysis were all largely stable between matched fresh and cryopreserved swabs taken from the same individual. Importantly, no “new” cell types (from healthy participants) were recovered from the freshly processed samples.

We interrogated each cell type for the expression of host factors utilized by common respiratory viruses to facilitate cellular entry (**Figure 1I**)^28,69–73^. We find *ACE2* expression highest among secretory cells and goblet cells, and to a lesser extent on ciliated cells, developing ciliated cells, deuterosomal cells, and squamous cells – suggesting these cells are likely targets for SARS-CoV-2 (and other betacoronaviruses that use ACE2 as their primary cellular entry factor). SARS-CoV-2 spike protein requires “priming” by host proteases such as *TMPRSS2, TMPRSS4, CTSL,* and *FURIN* for effective cell entry^69^. *TMPRSS2*, likely the principal host factor for SARS-CoV-2 S cleavage, is found in highest abundance in squamous cells, followed by modest expression in all other epithelial cell types. Similarly, *CTSL* (and other cathepsins) is found across diverse epithelial and myeloid cell types. *ANPEP* and *DPP4*, host receptors targeted by other human coronaviruses causing upper respiratory diseases, are found primarily in goblet cells and secretory cells^74,75^. As expected, *CDHR3*, the receptor utilized by Rhinovirus C, is found primarily in ciliated cells and developing ciliated cells^76^.

In order to assess compositional differences by disease severity, we grouped both SARS-CoV-2 positive and SARS-CoV-2 negative participants by their level of respiratory support according to the WHO scoring system: Control WHO 0 (comprising healthy SARS-CoV-2 PCR negative participants, n = 15), Control WHO 7-8 (SARS-CoV-2 PCR negative, incubated participants treated in the ICU for non-COVID-19 diagnoses, n = 6), COVID-19 WHO 1-5 (SARS-CoV-2 PCR positive, mild/moderate disease, n = 14), and COVID-19 WHO 6-8 (SARS-CoV-2 PCR positive, intubated, severe disease, n = 21). We compared proportional cell type abundances across these four groups (**Figure 1J-1N**). We found that the abundance of ciliated cells is significantly impacted by group (Kruskal-Wallis test with Dunn’s post-hoc testing, FDR-corrected p = 0.025), and is significantly reduced among COVID-19 WHO 6-8 participants compared to healthy controls (17.1 ± 3.6% (mean ± SEM) of COVID-19 WHO 6-8 samples are ciliated cells, compared to 46.7 ± 7.4% of Control WHO 0, p < 0.01) (**Figure 1J**). Deuterosomal cells, which represent a developmental intermediate as secretory/goblet cells differentiate into ciliated cells, are significantly increased among samples obtained from Control WHO 7-8, COVID-19 WHO 1-5, and COVID-19 WHO 6-8 samples, with the strongest increases observed among samples obtained from participants with severe COVID-19 compared to Control WHO 0 (**Figure 1L**). Likewise, developing ciliated cells are significantly increased among participants with severe COVID-19 (**Figure 1K**). The percentage of secretory cells is also dramatically increased among all COVID-19 participants compared to both the WHO 0 and WHO 7-8 control groups – 20.4 ± 5.0% (mean ± SEM) of all epithelial cells are secretory cells within severe COVID-19 participants, while mild/moderate COVID-19 participants contained 8.3 ± 2.8% secretory cells, and on average, fewer than 4% of cells per participant are secretory among either Control WHO 0 and Control WHO 7-8 samples (**Figure 1M**). The average percentage of goblet cells is higher in both groups of participants with COVID-19 compared to controls, but this difference does reach significance (**Figure 1N**). Intriguingly, expansion of secretory cells and loss of ciliated cells results in a net gain in epithelial diversity, calculated by Simpson’s index to calculate the richness of the epithelial “ecosystem” (**Figure 1O**).

### Epithelial Diversity and Remodeling Following SARS-CoV-2 Infection

Next, we sought to more completely delineate the diversity of epithelial cells through iterative clustering and sub-clustering among epithelial cell types (see **Methods**). This enabled us to divide the 10 “coarse” epithelial cell types into 25 “detailed” cell types/states (**Figures 2A-2E, Supplementary Figure 2A,** full differentiating gene lists for epithelial subtypes found in **Supplementary Table 1**). Among some cell types, we did not find additional within-type diversity, and thus the “coarse” annotations (**Figure 2A**) are equivalent to the “detailed” identities (**Figure 2D**). This applied to ionocytes, deuterosomal cells, developing secretory and goblet cells, basal cells, mitotic basal cells, and developing ciliated cells. We split goblet cells into 4 distinct detailed subtypes, each named by a representative defining marker or marker set. Likewise, secretory cells, squamous cells, and ciliated cells were all divided into multiple specialized subtypes. Some cellular subsets are similar to previously-described entities – including “*KRT24*^high^*KRT13*^high^ secretory cells”, which are highly similar to KRT13+ “hillock” cells, thought to be involved in airway epithelial responses to remodeling and inflammatory challenge^65,66^. Further, some cell types are defined by canonical cellular activation pathways, such as “interferon responsive” genes (e.g., *IFITM3*, *IFI6*, *MX1*) or “early response” factors (e.g., *JUN*, *EGR1*, *FOS*). Finally, some cell types contain specialized transcriptomic profiles, which, to our knowledge, have not been previously characterized. This included a subset of squamous cells expressing markers classically associated with vascular endothelial cells including *VWF* and *VEGFA*, as well as secretory populations expressing high abundances of multiple inflammatory cytokines, such as “*BPIFA1*^high^chemokine^high^ secretory cells” (chemokines include *CXCL8*, *CCL2*, *CXCL1*, and *CXCL3*) (**Figures 2D, 2E**).

**Figure 2.**
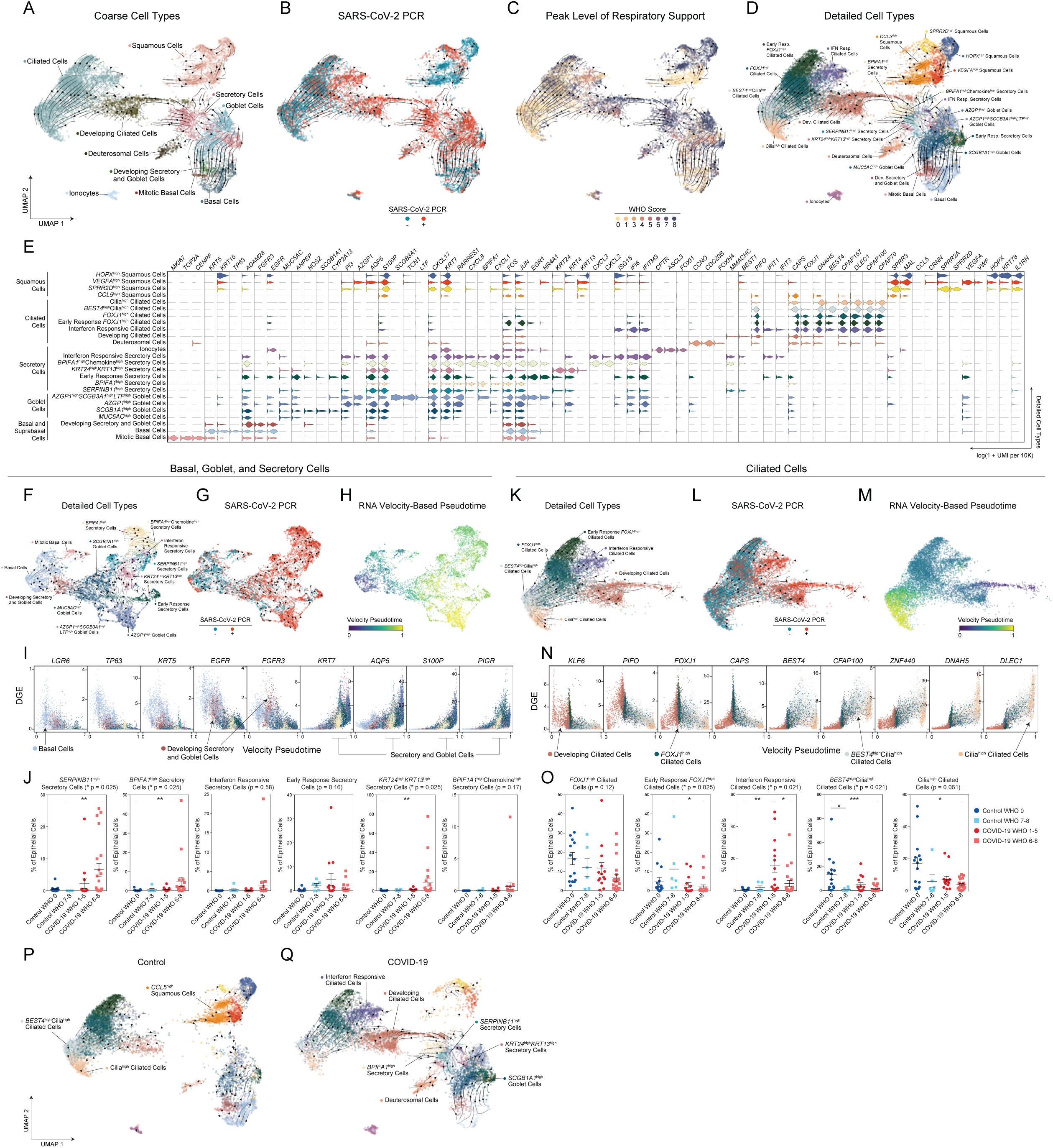
Altered Epithelial Cell Composition in the Nasopharynx During COVID-19. **A.** UMAP of 28,948 epithelial cell types following re-clustering, colored by coarse cell types. Arrows represent smoothed estimate of cellular differentiation trajectories **B.** UMAP as in **A**, colored by SARS-CoV-2 diagnostic PCR status. **C.** UMAP as in **A**, colored by peak level of respiratory support (WHO illness severity scale). **D.** UMAP as in **A**, colored by detailed cell types resolved by iterative re-clustering. **E.** Violin plots (log(1+normalized UMI per 10k)) of marker genes (FDR < 0.01) for detailed epithelial cell type annotations (as in **D**). **F.** UMAP of 9,209 basal, goblet, and secretory cells, following sub-clustering and resolution of detailed cell annotations. **G.** UMAP of only basal, goblet, and secretory cells as in **F**, colored by SARS-CoV-2 diagnostic PCR status. **H.** UMAP of only basal, goblet, and secretory cells as in **F**, colored by inferred velocity pseudotime (darker blue shades: precursor cells, intense yellow shades: more terminally differentiated cell types) **I.** Plot of gene expression by basal, goblet, and secretory cell velocity pseudotime for select genes (all significantly correlated with velocity expression). Points colored by detailed cell type annotations. **J.** Proportion of secretory cell subtypes (detailed annotation) by sample, normalized to all epithelial cells. Statistical test above graph represents Kruskal-Wallis test results across all groups (following FDR correction). Statistical significance asterisks within box represent results from Dunn’s post-hoc testing. * p < 0.05, ** p < 0.01, *** p < 0.001. Lines represent mean and S.E.M. **K.** UMAP of 13,913 ciliated cells, following sub-clustering and resolution of detailed cell annotations. **L.** UMAP of ciliated cells as in **K**, colored by SARS-CoV-2 PCR status at time of swab. **M.** UMAP of ciliated cells as in **K**, colored by inferred velocity pseudotime (darker blue shades: precursor cells, intense yellow shades: more terminally differentiated cell types). **N.** Plot of gene expression by ciliated cell velocity pseudotime for select genes (all significantly correlated with velocity expression). Points colored by detailed cell type annotations. **O.** Proportion of ciliated cell subtypes (detailed annotation) by sample, normalized to all epithelial cells. **P.** UMAP of 13,210 epithelial cells (using UMAP embedding from **A**) from SARS-CoV-2 PCR negative participants (Control). Arrows represent smoothed estimate of cellular differentiation trajectories (via RNA velocity) calculated using only cells from Control participants. **Q.** UMAP of 15,738 epithelial cells (using UMAP embedding from **A**) from SARS-CoV-2 PCR positive participants (COVID-19). Arrows represent smoothed estimate of cellular differentiation trajectories (via RNA velocity) calculated using only cells from COVID-19 participants. Named cell types highlight those significantly altered between disease groups. *See also Figure S2, Table S1*

We again examined the epithelial subtypes for their expression of host entry factors which facilitate viral entry among common upper respiratory pathogens (**Supplementary Figure 2B**). Here, we observed substantial within-cell type heterogeneity in *ACE2* expression among each of these cell types. Notably, among goblet cells, *AZGP1*^high^ goblet cells express the highest abundance of *ACE2* mRNA, suggesting this cell type may be a preferential target for SARS-CoV-2 infection. Likewise, early response secretory cells, *KRT24*^high^*KRT13*^high^ secretory cells, and interferon responsive secretory cells, all express elevated abundances of *ACE2*. Many other secretory and goblet cell types express detectable *ACE2*, but lower levels. Similarly, multiple detailed subsets of ciliated cells express *ACE2*, however cilia^high^ and *BEST4*^high^cilia^high^ ciliated cells did not appear to contain detectable levels of *ACE2* mRNA.

To map the differentiation and lineage relationships between epithelial cell types, we applied single-cell RNA velocity (scVelo), which leverages RNA splicing dynamics to infer developmental trajectories (**Methods**^77,78^). Overlaid on the UMAPs of cell type identities and associated metadata in **Figures 2A-2D**, vector fields (black lines and arrows) represent a smoothed estimate of cellular transitions based on RNA velocity. Globally, RNA velocity appropriately places basal cells and mitotic basal cells as the “root” or “origin” of cellular transitions, which then progresses through the developing secretory and goblet cells to the secretory cells and goblet cells. Developing ciliated cells and ciliated cells are placed “later” in the differentiation trajectory, distal to development of both secretory and deuterosomal cells, which is consistent with current models where ciliated cells represent a terminally differentiated state and may arise from these precursor cell types^64^. Together, this analysis enables us to map the developmental relationships between major epithelial cell compartments discussed above and connect the loss of “terminally differentiated” or “mature” cell types in COVID-19, e.g., ciliated cells, with the concurrent expansion of their apparent precursors: secretory, deuterosomal, and developing ciliated cells (**Figures 1J-1N, Supplementary Figure 2C**).

We next analyzed developmental transitions *among* detailed epithelial cell subtypes (annotated in **Figure 2D**) to better trace the relationships between finer-resolved subsets, and map alterations in cellular behavior and development during COVID-19. When considering only basal, goblet, and secretory cell subtypes, we found *LGR5, TP63*, *EGFR*, and *KRT5* expression gradually decline across basal and developing secretory and goblet cells, while expression of secretory and goblet cell specific markers such as *KRT7* and *AQP5* progressively increase (**Figure 2F-2I**). The majority of secretory and goblet clusters are represented by cells from individuals with positive SARS-CoV-2 PCR (as observed previously, **Figure 1K, 2G**), with significant expansion of *SERPINB11*^high^ secretory cells (which represent a “generic” or un-differentiated secretory subtype), *BPIFA1*^high^ secretory cells, and *KRT24*^high^*KRT13*^high^ secretory cells (which resemble KRT13+ “hillock” cells) among cells from individuals with severe COVID-19 (**Figure 2J**). Notably, transitions between detailed secretory and detailed goblet cells are more complex than those among the coarse cell types or as seen in ciliated cell subsets (discussed below). RNA velocity curves predict multiple routes for development between different secretory and goblet subtypes (**Figure 2F**), which suggests maintained capacity for differentiation and de-differentiation even among this “mature” cell type, and is consistent with the current understanding of respiratory secretory cell plasticity^79^.

Ciliated cell subtypes were analyzed by their RNA velocity and pseudotemporal ordering in the same manner (**Figures 2K-2N**). The velocity pseudotime predicts progression from developing ciliated cells, to *FOXJ1*^high^ ciliated cells, to *BEST4*^high^cilia^high^ ciliated cells, and terminating in cilia^high^ ciliated cells. (**Figure 2M**). Interferon responsive ciliated cells and early response *FOXJ1*^high^ ciliated cells represent phenotypic deviations from this ordered progression, and therefore appear collapsed/unresolved along this trajectory with the same pseudotime range as *FOXJ1*^high^ ciliated cells. Among COVID-19 participants, we observe decreased proportions of both cilia^high^ and *BEST4*^high^cilia^high^ ciliated cells, two cell subsets which represent the most terminally differentiated ciliated cell subtypes (**Figure 2O**). This effect is particularly pronounced among individuals with severe disease and suggests that the overall reduction in upper airway ciliated cells during COVID-19 (**Figure 1J)** preferentially affects terminally differentiated subsets, potentially due to delayed replenishing from secretory/deuterosomal precursors or enhanced susceptibility to viral-mediated pathogenesis. Among individuals with mild/moderate COVID-19, we find a substantial increase in the proportion of interferon responsive ciliated cells – averaging 15.9% of all epithelial cells among mild/moderate COVID-19 participants – compared to < 1% among healthy controls (**Figure 2O**).

Finally, we directly mapped the developmental transitions among nasal epithelial cells within Control (**Figure 2P**) or COVID-19 participants only (**Figure 2Q**). Confirming our above analysis, cells from Control participants poorly populated the intermediate regions that bridge secretory and goblet cell types to mature ciliated cells. Conversely, regions annotated as multiple secretory cell subsets and developing ciliated cells are uniquely captured from COVID-19 participants. Together, our analysis defines both the cellular diversity among cells collected from nasopharyngeal swabs, as well as the nuanced developmental relationships between epithelial cells of the upper airway. Further, we observe substantial expansion of immature/intermediate and specialized subtypes of secretory, goblet, and ciliated cells during COVID-19, presumably as a result of direct viral targeting and pathology, as well as part of the intrinsic capacity of the nasal epithelium to regenerate and repopulate following damage.

### Alterations to Nasal Mucosal Immune Populations in COVID-19

As with epithelial cells, we further clustered and annotated detailed immune cell populations. Multiple cell types could not be further subdivided from their coarse annotation (**Figure 1B, Supplementary Figure 3A-3E**), including mast cells, plasmacytoid DCs, B cells, and dendritic cells. Among macrophages (coarse annotation), we resolved 5 distinct subtypes (**Supplementary Figure 3B**). *FFAR4*^high^ macrophages are defined by expression of *FFAR4*, *MRC1*, *CHIT1*, and *SIGLEC11*, as well as chemotactic factors including *CCL18*, *CCL15*, genes involved in leukotriene synthesis (*ALOX5, ALOX5AP, LTA4H*), and toll-like receptors *TLR8* and *TLR2* (**Supplementary Figure 3F**, full differentiating gene lists for immune subtypes found in **Supplementary Table 1**). Interferon responsive macrophages are distinguished by elevated expression of anti-viral genes such as *IFIT3, IFIT2, ISG15,* and *MX1*, akin to the epithelial subsets labeled “interferon responsive”, along with *CXCL9*, *CXCL10*, *CXCL11*, which are likely indicative of IFNγ stimulation. *MSR1*^high^*C1QB*^high^ macrophages are defined by cathepsin expression (*CTSD, CTSL, CTSB*) and elevated expression of complement (*C1QB, C1QA, C1QC*), and lipid binding proteins (*APOE, APOC,* and *NPC2*). The fourth “specialized” subtype of macrophage we term “inflammatory macrophages”, which uniquely express inflammatory cytokines such as *CCL3, CCL3L1, IL1B, CXCL2,* and *CXCL3*. The remaining “*ITGAX*^high^” macrophages are distinguished from other immune cell types by *ITGAX*, *VCAN, PSAP, FTL, FTH1* and *CD163* (though these genes are shared by other specialized macrophages subsets). T cells are largely *CD69* and *CD8A* positive, consistent with a T resident memory-like phenotype^80^, and we are not able to resolve a separate cluster of CD4 T cells. Two specialized subtypes of CD8 T cells are annotated from this dataset: one defined by exceptionally high expression of early response genes (*FOSB, NR4A2,* and *CCL5*), and the other termed “interferon responsive cytotoxic CD8 T cells”, defined by granzyme and perforin expression (*GZMB, GZMA, GNLY, PRF1, GZMH*), anti-viral genes (*ISG20, IFIT3, APOBEC3C, GBP5*) and genes associated with effector CD8 T cell function (*LAG3, IL2RB, IKZF3, TBX21*).

Among immune cells, macrophages appear to markedly increase in abundance relative to other immune cell types during severe COVID-19 (**Supplementary Figure 3G, 3H**). Multiple specialized myeloid cell types are uniquely detected and enriched among COVID-19 participants, albeit in a subset of participants, and biased to severe COVID-19 cases: *ITGAX*^high^ macrophages, *FFAR*4^high^ macrophages, inflammatory macrophages, and interferon responsive macrophages (**Supplementary Figure 3H**). Through rare, plasmacytoid DCs and mast cells are only recovered as > 1% of immune cells among COVID-19 participants, though this enrichment did not reach statistical significance. Somewhat surprisingly, we do not find T cells and T cell subtypes to be dramatically altered between disease groups. Finally, we assessed the correlation between distinct cell types across all participants. When samples from all disease groups were considered, we find that proportional abundance of dendritic cells, mast cells, and macrophages were highly-correlated with one another (p < 0.01), likely indicative of the coordinated recruitment of these immune subtypes during inflammation. Among detailed immune cell types, interferon responsive macrophages are correlated with interferon responsive cytotoxic CD8 T cells (p < 0.01, **Supplementary Figure 3I**), suggesting potential direct communication between *IFNG*-expressing tissue resident T cells and *CXCL9/10/11* expressing myeloid cells.

Collectively, these analyses demonstrate how the epithelial and immune compartments are dramatically altered during COVID-19, likely reflecting both protective anti-viral and regenerative responses, as well as pathologic changes underlying progression to severe disease.

### Cellular Behaviors Associated with COVID-19 Severity

Thus far, we have characterized how severe COVID-19 elicits major cell compositional changes within the nasopharyngeal mucosa, including expansion of the secretory cell/deuterosomal cell compartments associated with lost mature ciliated cells, and recruitment of highly inflammatory myeloid cells. Next, we examined how each individual cell type responds across the full spectrum of disease severity. Here, we analyzed pairwise comparisons between Control WHO 0, COVID-19 WHO 1-5 (mild/moderate), and COVID-19 WHO 6-8 (severe), and compared both high-level “coarse” cell types, and “detailed” cell subsets (**Figure 3A, Supplementary Figure 4A, Supplementary Tables 2-4**). Among all coarse cell types, the largest magnitude transcriptional changes (measured by the number of differentially expressed (DE) genes with FDR < 0.001, and log fold change > 0.25) are observed primarily within the epithelial compartment, most strikingly within ciliated cells, developing ciliated cells, secretory cells, goblet cells, and ionocytes (**Supplementary Figure 4A**). Among detailed cell types, we observed the largest transcriptional changes among *AZGP1*^high^ goblet cells, early response *FOXJ1*^high^ ciliated cells, *FOXJ1*^high^ ciliated cells, *MUC5AC*^high^ goblet cells, *SERPINB11*^high^ secretory cells, early response secretory cells, and interferon responsive ciliated cells. Broadly, major differences are observed in the *identity* of cell types with large transcriptional responses – with mild/moderate COVID-19 driving differences principally in *MUC5AC*^high^ goblet cells and ionocytes, while severe COVID-19 included major perturbations among basal cells and *AZGP1*^high^ goblet cells. Ciliated subsets are profoundly altered in both mild/moderate and severe COVID-19 compared to cells from Control WHO 0 participants. Finally, when we directly compared mild/moderate to severe COVID-19, we found that multiple cell types show robust transcriptional changes, most drastically among ciliated cell subtypes (interferon responsive ciliated cells, *FOXJ1*^high^ ciliated cells, early response *FOXJ1*^high^ ciliated cells, developing ciliated cells), ionocytes, *SERPINB11*^high^ secretory cells, early response secretory cells, and *AZGP1*^high^ goblet cells.

**Figure 3.**
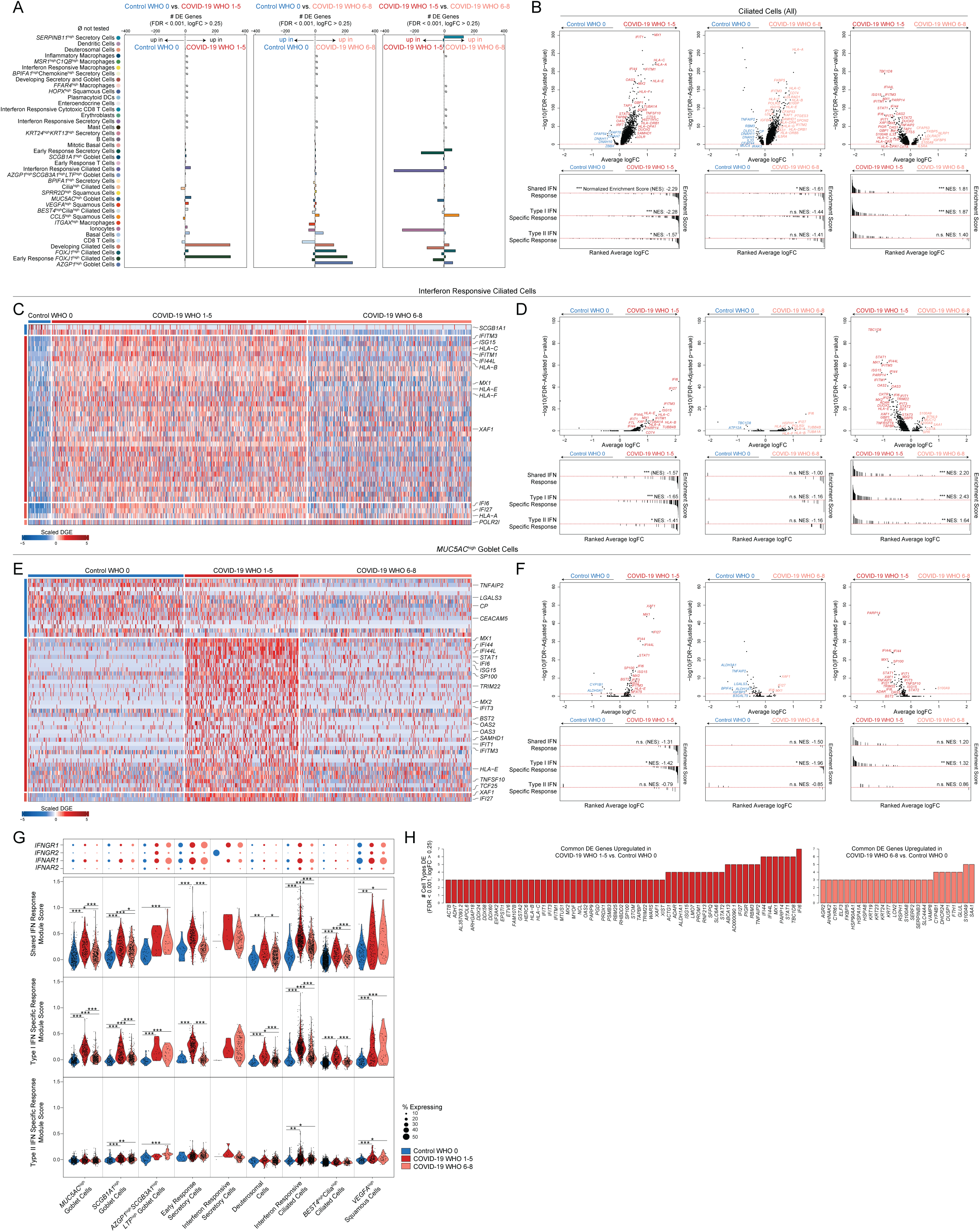
Cell-Type Specific and Shared Transcriptional Responses During COVID-19. **A.** Abundance of significantly differentially expressed (DE) genes by detailed cell types between Control WHO 0 vs. COVID-19 WHO 1-5 samples (left), Control WHO 0 and COVID-19 WHO 6-8 samples (middle), COVID-19 WHO 1-5 and COVID-19 WHO 6-8 samples (right). Restricted to genes with FDR-corrected p < 0.001, log2 fold change > 0.25 (likelihood ratio test assuming an underlying negative binomial distribution). ø = comparison not tested, too few cells (< 10). **B.** Top: Volcano plots of average log fold change (FC) vs. −log_10_(FDR-adjusted p-value) for ciliated cells (all, coarse annotation). Left: Control WHO 0 vs. COVID-19 WHO 1-5 (mild/moderate). Middle: Control WHO 0 vs. COVID-19 WHO 6-8 (severe). Right: COVID-19 WHO 1-5 (mild/moderate) vs. COVID-19 WHO 6-8 (severe). Horizontal red dashed line: FDR-adjusted p-value = 0.05. Bottom: gene set enrichment analysis plots across shared, type I interferon specific, and type II interferon specific stimulated genes. Genes ranked by their average log FC between each comparison. Black lines represent the ranked location of genes belonging to the annotated gene set. Bar height represents running enrichment score (NES: Normalized Enrichment Score). P-values following Bonferroni-correction: * p < 0.05, ** p < 0.01, *** p < 0.001. **C.** Heatmap of significantly DE genes between interferon responsive ciliated cells from different disease groups. Values represent row(gene)-scaled digital gene expression (DGE) following log(1+UMI per 10K) normalization. **D.** Top: Volcano plots related to **C**. Average log FC vs. −log_10_(FDR-adjusted p-value) for interferon responsive ciliated cells. Horizontal red dashed line: FDR-adjusted p-value = 0.05. Bottom: gene set enrichment analysis across shared, type I, and type II interferon stimulated genes. **E.** Heatmap of significantly DE genes between *MUC5AC*^high^ goblet cells from different disease groups. Values represent row(gene)-scaled digital gene expression (DGE) following log(1+UMI per 10K) normalization. **F.** Top: Volcano plots related to **E**. Average log FC vs. −log_10_(FDR-adjusted p-value) for *MUC5AC*^high^ goblet cells. Horizontal red dashed line: FDR-adjusted p-value = 0.05. Bottom: gene set enrichment analysis across shared, type I, and type II interferon stimulated genes. **G.** Top: Dot plot of *IFNGR1, IFNGR2*, *IFNAR1, and IFNAR2* gene expression among subset of detailed epithelial subtypes. Bottom: Violin plots of module scores, split by Control WHO 0 (blue), COVID-19 WHO 1-5 (red), and COVID-19 WHO 6-8 (pink). Gene modules represent transcriptional responses of human basal cells from the nasal epithelium following *in vitro* treatment with IFNα or IFNγ. Significance by Wilcoxon signed-rank test. P-values following Bonferroni-correction: * p< 0.05, ** p < 0.01, *** p < 0.001. **H.** Common DE genes across detailed cell types. Left (red): genes upregulated in multiple cell types when comparing COVID-19 WHO 1-5 vs. Control WHO 0. Right (pink): genes upregulated in multiple cell types when comparing COVID-19 WHO 6-8 vs. Control WHO 0. *See also Figures S3, S4, Tables S2-S4*

We further examined the specific DE genes among ciliated cells (all, coarse annotation) between each group (**Figure 3B, Supplementary Tables 2-4**). Compared to ciliated cells from Control WHO 0 participants, cells from both mild/moderate COVID-19 and severe COVID-19 robustly upregulate genes involved in the host response to virus, including *IFI27, IFIT1, IFI6, IFITM3,* and *GBP3*, and both groups induce expression of MHC-I and MHC-II genes (including *HLA-A, HLA-C, HLA-F, HLA-E, HLA-DRB1, HLA-DRA*) and other factors involved in antigen processing and presentation (**Supplementary Figures 4B, 4C**). Notably, large sets of interferon-responsive and anti-viral genes are exclusively induced among ciliated cells from COVID-19 WHO 1-5 participants when compared to Control WHO 0 participants. In a direct comparison of ciliated cells from mild/moderate to severe COVID-19, the cells from individuals with mild/moderate disease show strong upregulation of diverse anti-viral factors, including *IFI44L, STAT1, IFITM1, MX1, IFITM3, OAS1, OAS2, OAS3, STAT2, TAP1, HLA-C, ADAR, XAF1, IRF1, CTSS, CTSB*, and many others (**Supplementary Figure 4C**). Ciliated cells from severe COVID-19 uniquely upregulate *IL5RA* and *NLRP1* (compared to both control and mild/moderate COVID-19). Together, these DE gene sets are suggestive of exposure to secreted inflammatory factors and type I/II/III interferons, as well as direct cellular sensing of viral products. Using previously published data from human nasal basal cells treated *in vitro* with either type I (IFNα) or type II (IFNγ) interferon^29^, we created gene sets that represent the “shared” gene responses to type I and type II interferon, and the cellular responses specific to either type (**Figure 3B**). Using gene set enrichment analysis, we tested whether the genes that discriminate ciliated cells from different groups (e.g., mild/moderate COVID-19 vs. severe COVID-19) imply exposure to specific interferon types. We found that ciliated cells in mild/moderate COVID-19 robustly induce type I interferon-specific gene signatures, both compared to cells from healthy controls, as well cells from severe COVID-19. Further, when compared to cells from healthy individuals, ciliated cells from individuals with severe COVID-19 did not significantly induce type I or type II interferon responsive genes, potentially underlying poor control of viral spread.

We next investigated whether these effects were observed among other cell types and subsets. Surprisingly, even *among* cells defined as “interferon responsive” ciliated cells, cells from mild/moderate COVID-19 participants express higher fold changes of interferon-responsive genes compared to cells from COVID-19 WHO 6-8 participants or Control WHO 0 (**Figures 3C, 3D, Supplementary Tables 2-4**). Other detailed epithelial and immune cell types display a similar pattern: broad interferon-responsive genes (largely type I specific) are strongly upregulated among cells from mild/moderate COVID-19 participants, while cells from severe COVID-19 upregulate few shared markers with mild/moderate COVID-19 participants, and instead skew towards inflammatory genes such as *S100A8* and *S100A9* instead of anti-viral factors (**Figures 3E-3H, Supplementary Figures 3J-3L, 4D**). In some cases, cells from individuals with severe COVID-19 express levels of interferon responsive or anti-viral genes indistinguishable from healthy controls. Strongest induction of type I specific interferon responses among mild/moderate COVID-19 cases is observed in *MUC5AC*^high^ goblet cells, *SCGB1A1*^high^ goblet cells, early response secretory cells, deuterosomal cells, interferon responsive ciliated cells, and *BEST4*^high^cilia^high^ ciliated cells (**Figure 3G**). Rare cell types from individuals with severe COVID-19 induce comparable type I interferon responses to their mild/moderate counterparts, including *AZGP1*^high^*SCGB3A1*^high^*LTF*^high^ goblet cells, interferon responsive secretory cells, and *VEGFA*^high^ squamous cells. Expression of type II interferon specific genes is globally blunted across all cell types from COVID-19 samples when compared to type I interferon module scores (**Figure 3G, Supplementary Figures 3K, 4D**). Further, the absence of a transcriptional response to secreted interferon cannot be explained by a lack of either interferon alpha receptor (*IFNAR1, IFNAR2*) or interferon gamma receptor (*IFNGR1, IFNGR2*) expression. Previous work has identified *ACE2*, the host receptor for SARS-CoV-2, as among the interferon-induced genes in nasal epithelial cells, with uncertain significance for SARS-CoV-2 infection^29,81–83^. Indeed, we find modest upregulation of this gene among cells from COVID-19 participants compared to healthy controls. Further, some of the cell subtypes identified as expanded during COVID-19 (e.g., interferon responsive ciliated cells, *BPIFA1*^high^ secretory cells, *BPIFA1*^high^chemokine^high^ secretory cells, and *KRT24*^high^*KRT13*^high^ secretory cells) express relatively high abundances of *ACE2* (**Supplementary Figure 4E**).

A sizable proportion of COVID-19 participants in our study were concurrently treated with corticosteroids, which mediate broad anti-inflammatory and immunosuppressive effects. We were curious as to whether depressed interferon and anti-viral responses could be explained by higher rates of corticosteroid treatment among the severe COVID-19 group (**Table 1**). We therefore stratified our groups further into Steroid-Treated vs. Untreated and assessed expression of genes previously identified as DE between Control WHO 0, COVID-19 WHO 1-5, and COVID-19 WHO 6-8. For some genes, corticosteroid treatment is associated with a partially suppressed interferon response *within* each group – for instance, ciliated cells from Untreated COVID-19 WHO 1-5 participants show higher abundances of *IFITM1, OAS2, IFI6*, and *IFI27* than their Steroid-Treated counterparts – while still maintaining strong differences in expression *between* groups (with abundance in COVID-19 WHO 1-5 > COVID-19 WHO 6-8 > COVID-19 WHO 0, see annotations on **Supplementary Figure 4C**). Interestingly, induction of *FKBP5* expression among ciliated cells from severe COVID-19 participants is fully explained by corticosteroid treatment, which is consistent with the role for this protein in modulating glucocorticoid receptor activity. Other sets of anti-viral genes were equivalently expressed within each group, independent of treatment, including *STAT1*, *STAT2, IFI44,* and *ISG15*. For many anti-viral factors in multiple cell types, we observed no effect of corticosteroid treatment on the intrinsic anti-viral response during COVID-19.

Together, these data demonstrate global blunting of the anti-viral/interferon response among nasopharyngeal epithelial cells during severe COVID-19. Individuals with mild/moderate COVID-19 recurrently upregulate interferon-responsive factors including *STAT1, MX1, HLA-B, HLA-C*, among others (compared to matched cell types among Control WHO 0 participants), while cells from individuals with severe COVID-19 repeatedly induced a distinct set of genes, including *S100A9, S100A8* and stress response factors (*HSPA8, HSPA1A, DUSP1,* **Figure 3H**). We next attempted to query the source of local interferon, particularly in the COVID-19 WHO 1-5 samples where cell types appeared to be maximally responding to interferon stimulation. Notably, we expect many of the tissue-resident immune cells to reside principally within the deeper lamina propria and submucosal spaces, and are therefore poorly represented in our dataset due to sampling type (swabbing of surface epithelial cells)^65^. Accordingly, we find exceedingly few immune cell types producing interferons: *IFNA* and *IFNB* are absent, rare *IFNL1* UMI are observed among T cells and macrophages, and *IFNG* is robustly produced from interferon responsive cytotoxic CD8 T cells, despite limited evidence for type II responses among epithelial cells (**Supplementary Figure 4F**). Further, we could not detect expression of any interferon types among epithelial cells, which is differs dramatically from previous observations of robust type I/III interferon expression among nasal ciliated cells during influenza A and B infection (also captured via Seq-Well S^3 84^ (**Supplementary Figure 4F**). Rather, we observe robust induction of other inflammatory molecules from both immune and epithelial cell types. *CXCL8* is produced by several specialized secretory cell types, including those uniquely expanded in COVID-19. Inflammatory macrophages and interferon responsive macrophages represent the primary sources of local *TNF*, *IL6*, and *IL10*, and uniquely express high abundances of chemoattractant molecules such as *CCL3, CCL2,* and *CXCL8.* Interestingly, interferon responsive macrophages appear to be a principal source of *CXCL9, CXCL10,* and *CXCL11* (**Supplementary Figures 4F**).

### Co-Detection of Viral and Host RNA and Correlates of Nasal Viral Load

Given a comprehensive picture of host cell biology during COVID-19 and across the spectrum of disease severity, we next tested whether the observed epithelial phenotypes were associated with altered local viral abundance. ScRNA-seq protocols utilize poly-adenylated RNA capture and reverse transcription to generate snapshots of the transcriptional status of each individual cell. As several pathogens and commensal microbes also utilize poly-adenylation for RNA intermediates, or contain poly-adenylated stretches of RNA within their genomes, they may also be represented within scRNA-seq libraries. First, to perform an unbiased search for co-detected viral, bacterial, and fungal genomic material, we used metatranscriptomic classification to assign reads according to a comprehensive reference database (previously described, see **Methods**)^85,86^. As expected, the majority (28/38) of swabs from individuals with COVID-19 contain reads classified as SARS coronavirus species (**Figure 4A, Supplementary Figures 5A-5C**). Among samples containing SARS coronavirus genomic material, the read abundance ranged from 2e0 to 8.8e6 reads (1.8e-3 to 1.9e4 reads/million (M) total reads). We found little evidence for co-occurring respiratory viruses, which may be partially explained by the season when many of the swabs were collected (April-September 2020) and concurrent social distancing practices. Swabs from two individuals contain rare reads classified as Influenza A virus species (maximum 5 reads per donor, within range for spurious classification), and we observe no evidence for other seasonal human coronaviruses, Influenza B virus, metapneumovirus, or orthopneumovirus. Swabs from two individuals with mild/moderate COVID-19 contain exceptionally high abundances of reads classified as Rhinovirus A (2.1e5 and 2.4e5 reads). Finally, we recover low abundances of SARS coronavirus assigned reads from two participants from the Control WHO 0 group.

**Figure 4.**
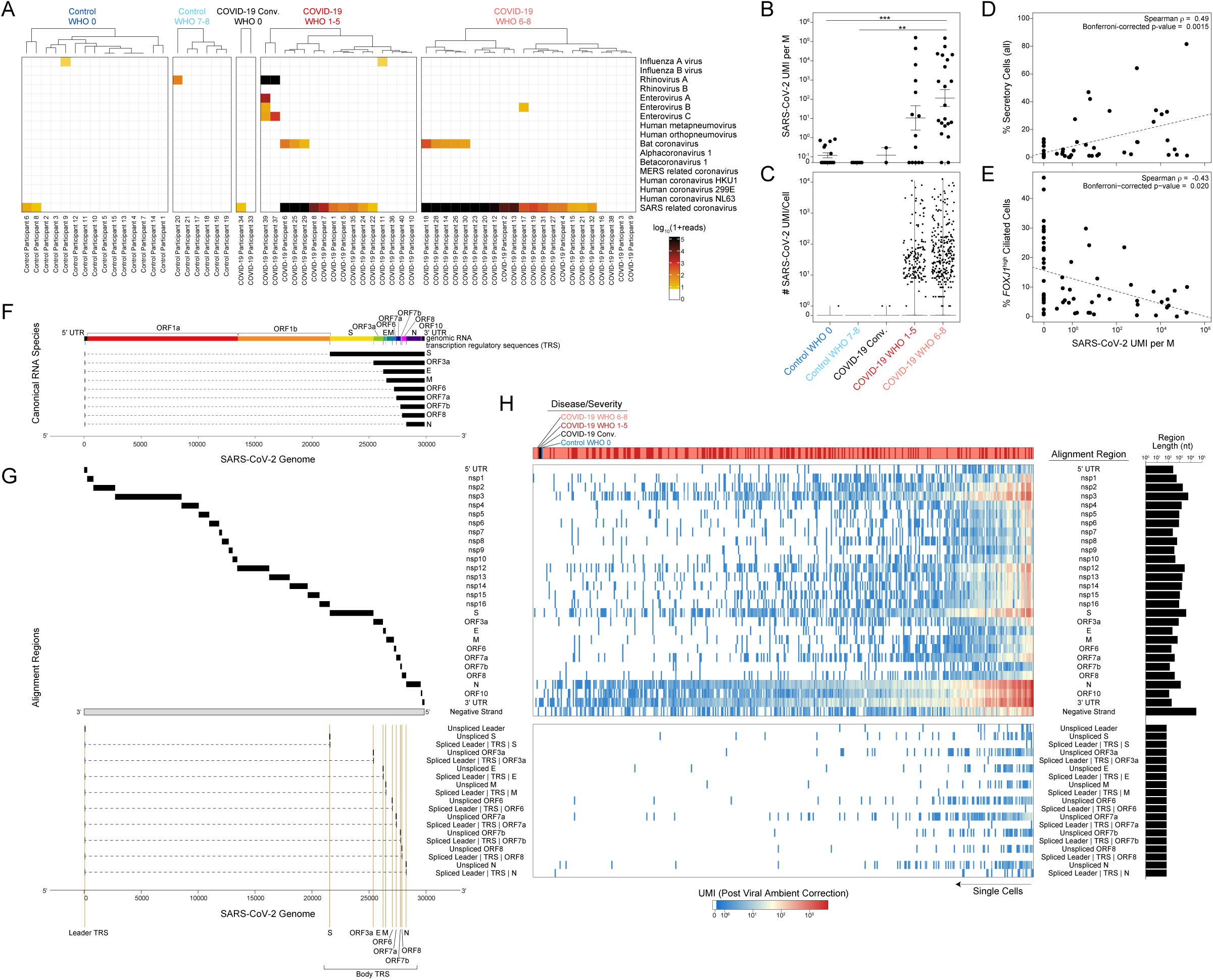
Co-Detection of Human and SARS-CoV-2 RNA. **A.** Metatranscriptomic classification of all scRNA-seq reads using Kraken2 (**Methods**). Results shown from selected respiratory viruses. Only results with > 5 reads are shown. **B.** Normalized abundance of SARS-CoV-2 aligning UMI from all scRNA-seq reads (including those derived from ambient cell barcodes). P < 0.0001 by Kruskal-Wallis test. Pairwise comparisons using Dunn’s post-hoc testing. ** p < 0.01, *** p < 0.001. **C.** SARS-CoV-2 UMI per high-complexity single-cell transcriptome. Results following correction for ambient viral reads. **D.** Proportional abundance of secretory cells (all, coarse annotation) vs. total SARS-CoV-2 UMI (normalized to M total UMI). **E.** Proportional abundance of *FOXJ1*^high^ ciliated cells vs. total SARS-CoV-2 UMI (normalized to M total UMI). **F.** Schematic for SARS-CoV-2 genome and subgenomic RNA species. **G.** Schematic for SARS-CoV-2 genomic features annotated in the custom reference genome. **H.** Heatmap of SARS-CoV-2 gene expression among SARS-CoV-2 RNA+ single cells (following correction for ambient viral reads). Top color bar indicates disease group (red: COVID-19 WHO 1-5, pink: COVID-19 WHO 6-8, black: COVID-19 Convalescent, blue: Control WHO 0). Middle heatmap: SARS-CoV-2 genes and regions organized from 5’ to 3’. Bottom heatmap: alignment to 70-mer regions directly surrounding viral transcription regulatory sequence (TRS) sites, suggestive of spliced RNA species (joining of the leader to body regions) vs. unspliced RNA species (alignment across TRS). *See also Figures S5, S6*

Next, we analyzed all SARS-CoV-2-aligned UMI following alignment to a joint genome containing both human and SARS-CoV-2^87^. We took the sum of all SARS-CoV-2 aligning UMI from a given participant – both associated with high-complexity single-cell transcriptomes and ambient RNA – as a representative measure of the total SARS-CoV-2 burden within the tissue microenvironment. As observed using metatranscriptomic classification, we found relatively low/spurious alignments to SARS-CoV-2 among Control participants, while swabs from COVID-19 participants contained a wide range of SARS-CoV-2 aligning reads (**Figure 4B, 4C, Supplementary Figures 5D, 5E**). Samples from participants with severe COVID-19 contained significantly higher abundances of SARS-CoV-2 aligning reads than both control groups, with an average of 1.1e2 ± 2.8e0 (geometric mean ± SEM) UMI per million (M) aligned UMI (ranging from 0 to 1.5e5 per sample). Swabs from participants with mild/moderate COVID-19 contained slightly fewer SARS-CoV-2 aligning UMI, with an average of 1.1e1 ± 4.3e0 (geometric mean ± SEM) UMI per M.

Given the large diversity in SARS-CoV-2 abundance across all COVID-19 participants, we interrogated whether cell composition correlated with total SARS-CoV-2 (**NB**: contemporaneous work by our group has evaluated the accuracy of single-cell RNA-seq-derived estimates of total SARS-CoV-2 abundance with more established protocols such as qRT-PCR)^88^. Among all cell types, we observe that secretory cells are significantly positively correlated with the total viral abundance (Spearman’s rho = 0.49, Bonferroni-corrected p = 0.0015), while *FOXJ1*^high^ ciliated cells are significantly negatively correlated (Spearman’s rho = −0.43, Bonferroni-corrected p = 0.020, **Figures 4D, 4E**). This observation is in line with findings outlined in **Figures 1** and **2** where epithelial pathology during SARS-CoV-2 infection manifests as loss of mature ciliated cell types, which may stimulate secretory cells to expand and repopulate lost epithelial cell types, although direct virally-mediated effects on secretory cell expansion have not been ruled out. Next, we binned the samples from COVID-19 participants into “Viral Low” and “Viral High” groupings (based on an arbitrary cutoff of 1e3 SARS-CoV-2 UMI per M; our findings (below) are robust to a range of partition choices, **Supplementary Figures 5E, 5F**). Interferon responsive ciliated cells are expanded among “Viral High” COVID-19 samples and plasmacytoid DCs are absent from “Viral High” samples. Finally, in a subset of patients for whom we obtained matched plasma samples on the same day of nasopharyngeal swab (n = 36), we observe SARS-CoV-2 UMI abundance is inversely correlated with the SARS-CoV-2 IgM and IgG titers (**Supplementary Figure 5G**), suggesting that individuals with high viral loads are sampled early in their disease trajectory^89–91^.

### Cellular Targets of SARS-CoV-2 Infection in the Nasopharynx

Next, we aimed to differentiate SARS-CoV-2 UMI derived from ambient or low-complexity cell barcodes from those likely reflecting intracellular RNA molecules within high-complexity single-cell transcriptomes^84,88,92,93^. First, we filtered to only viral UMIs associated with cells presented in **Figure 1**, thereby removing those associated with low-complexity or ambient-only cell barcodes (**Supplementary Figure 5H**). Using a combination of computational tools to 1) estimate the proportion of ambient RNA contamination per single cell and 2) estimate the abundance of SARS-CoV-2 RNA within the extracellular/ambient environment (i.e., not cell-associated) per sample, we were able to test whether the amount of viral RNA associated with a given single-cell transcriptome was significantly higher than would be expected from ambient spillover. Together, this enabled us to identify cell barcodes whose SARS-CoV-2 aligning UMI were likely driven by spurious contamination, and annotate single cells that contain probable cell-associated or intracellular SARS-CoV-2 RNA (**Figure 4C, Supplementary Figure 5H**). Across all single cells, we recover 415 high-confidence SARS-CoV-2 RNA+ cells across 21 participants, and we confirmed that cell assignment as “SARS-CoV-2 RNA+” is not driven by technical factors such as sequencing depth or cell complexity (**Supplementary Figure 5I**). 262 SARS-CoV-2 RNA+ cells are from participants with severe COVID-19 and 150 from mild/moderate COVID-19. We detect 3 SARS-CoV-2 RNA+ cells from participants with negative SARS-CoV-2 PCR: two from a participant identified as “COVID-19 Convalescent”, and one from a Control participant. Among participants with any SARS-CoV-2 RNA+ cell, we detect 20 ± 7 (mean ± SEM) SARS-CoV-2 RNA+ cells per sample (range 1-119), amounting to 4 ± 1.3% (range 0.1-24%) of the total recovered cells per sample. *Within* a given single cell, the abundance of SARS-CoV-2 UMI ranges from 1 to 12,612, corresponding to 0.01-98% of all human and viral UMI per cell.

To further understand the biological significance of SARS-CoV-2 aligning UMI within a single cell, and to better identify cells with the highest-likelihood of actively supporting viral replication, we analyzed the specific viral sequences and their alignment regions in the viral genome^87,94,95^. During SARS-CoV-2 infection, viral uncoating from endosomal vesicles releases the positive, single-stranded, 5’ capped, poly-adenylated genome into the host cytosol (**Figures 4F, 4G**). Here, translation of non-structural proteins proceeds first by templating directly off of the viral genome, generating a replication and transcription complex. The viral replication complex then produces both 1) negative strand genomic RNA intermediates, which serve as templates for further positive strand genomic RNA and 2) nested subgenomic mRNAs which are constructed from a 5’ leader sequence fused to a 3’ sequence encoding structural proteins for production of viral progeny (e.g., Spike, Envelope, Membrane, Nucleocapsid). Generation of nested subgenomic mRNAs relies on discontinuous transcription occurring between pairs of 6-mer transcriptional regulatory sequences (TRS), one 3’ to the leader sequence (termed leader TRS, or TRS-L), and others 5’ to each gene coding sequence (termed body TRS, or TRS-B)^96^. We reasoned that SARS-CoV-2 aligning UMI could be readily distinguished by their strandedness (aligning to the negative vs. positive strand) and whether they fell within coding regions, across intact TRS (indicating RNA splicing had not occurred for that RNA molecule at that splice site) or across a TRS with leader-to-body fusions (corresponding to subgenomic RNA, **Figure 4F, 4G, Supplementary Figure 6A**). Single cells containing higher abundances of spliced TRS or negative strand aligning reads are therefore more likely to represent truly virally-infected cells with a functional viral replication and transcription complex. Critically, the co-detection of host transcriptomic and viral genomic material associated with a single cell barcode cannot definitively establish the presence of intracellular virus and/or productive infection. Rather, below we integrate these and other aspects of the host and viral transcriptomes to refine and contextualize our confidence in “SARS-CoV-2 RNA+” cells.

The majority of SARS-CoV-2 aligning UMI among SARS-CoV-2 RNA+ cells are found heavily biased towards the 3’ end of the genome, attributed to the 3’ UTR, ORF10, and N gene regions, as expected due to poly-A priming (**Figure 4H**). A majority (68.7%) of SARS-CoV-2 RNA+ cells contain reads aligning to the viral negative strand, increasing the likelihood that many of these cells represent true targets of SARS-CoV-2 virions *in vivo*. In addition to negative strand alignment, we find roughly ∼ 1/4 of the SARS-CoV-2 RNA+ cells contain at least 100 UMI that map to more than 20 distinct viral genomic locations per cell. Finally, comparing spliced to unspliced UMI, we find a minor fraction of cells with reads mapping directly across a spliced TRS sequence (4.6%), while 35% of SARS-CoV-2 RNA+ cells contain reads mapping across the equivalent 70mer window around an unspliced TRS. Notably, single cells containing reads aligning to spliced (subgenomic) RNA are heavily skewed toward those cells that contain the highest overall abundances of viral UMI – this may be an accurate reflection of coronavirus biology, wherein subgenomic RNA are most frequent within cells robustly producing new virions and total viral genomic material, but also points to inherent limitations in the detection of low-frequency RNA species by single-cell RNA-seq technologies.

Next, we integrated 1) the strand and splice information among SARS-CoV-2 aligning UMIs, 2) participant-to-participant diversity and 3) cell type annotations to gain a comprehensive picture of the identity and range of SARS-CoV-2 RNA+ cells within the nasopharyngeal mucosa (**Figures 5A-D, Supplementary Figures 6A-6E**). We observe incredible diversity in both the identity of SARS-CoV-2 RNA+ cells, as well as the distribution of SARS-CoV-2 RNA+ cells within and across participants. The majority of SARS-CoV-2 RNA+ cells are ciliated, goblet, secretory, or squamous. Highest-confidence SARS-CoV-2 RNA+ cells (containing UMI aligning across a spliced TRS, negative strand UMI, and > 100 SARS-CoV-2 UMI/cell) tended to be found among *MUC5AC*^high^ goblet cells, *AZGP1*^high^ goblet cells, *BPIFA1*^high^ secretory cells, *KRT24*^high^*KRT13*^high^ secretory cells, *CCL5*^high^ squamous cells, developing ciliated cells, and each ciliated cell subtype. A high proportion of interferon responsive macrophages contained SARS-CoV-2 genomic material, and rare *ITGAX*^high^ macrophages are found to contain UMI aligning to viral negative strand or spliced TRS regions – likely representing myeloid cells that have recently engulfed virally-infected epithelial cells or free virions. We did not find major differences in the presumptive cellular tropism by COVID-19 severity. A few cell types are commonly found to be SARS-CoV-2 RNA+ across all participants (including participants with only rare viral RNA+ cells): most frequently, participants have at least one developing ciliated or squamous cell with SARS-CoV-2 RNA, followed by *MUC5AC*^high^ goblet cells, cilia^high^ ciliated cells, and *FOXJ1*^high^ ciliated cells (**Figure 5D**). However, among the individuals with the highest abundances of SARS-CoV-2 RNA+ cells, viral RNA is spread broadly across many different cell types, including those outside of the expected tropism for SARS-CoV-2 (e.g., also found within basal cells, ionocytes). Further, the cell types harboring the highest proportions of SARS-CoV-2 RNA+ cells represent the same cell types uniquely expanded or induced within COVID-19 participants, such as *KRT24*^high^*KRT13*^high^ secretory cells, *AZGP1*^high^ goblet cells, and interferon responsive ciliated cells, and contain the highest abundances of *ACE2*-expressing cells (**Figure 5C, Supplementary Figure 6F**). Whether these cell types represent specific phenotypes elicited by intrinsic viral infection (potentially alongside induction of anti-viral genes) or are uniquely susceptible to SARS-CoV-2 entry (e.g., enhanced entry factor expression) will require further investigation.

**Figure 5.**
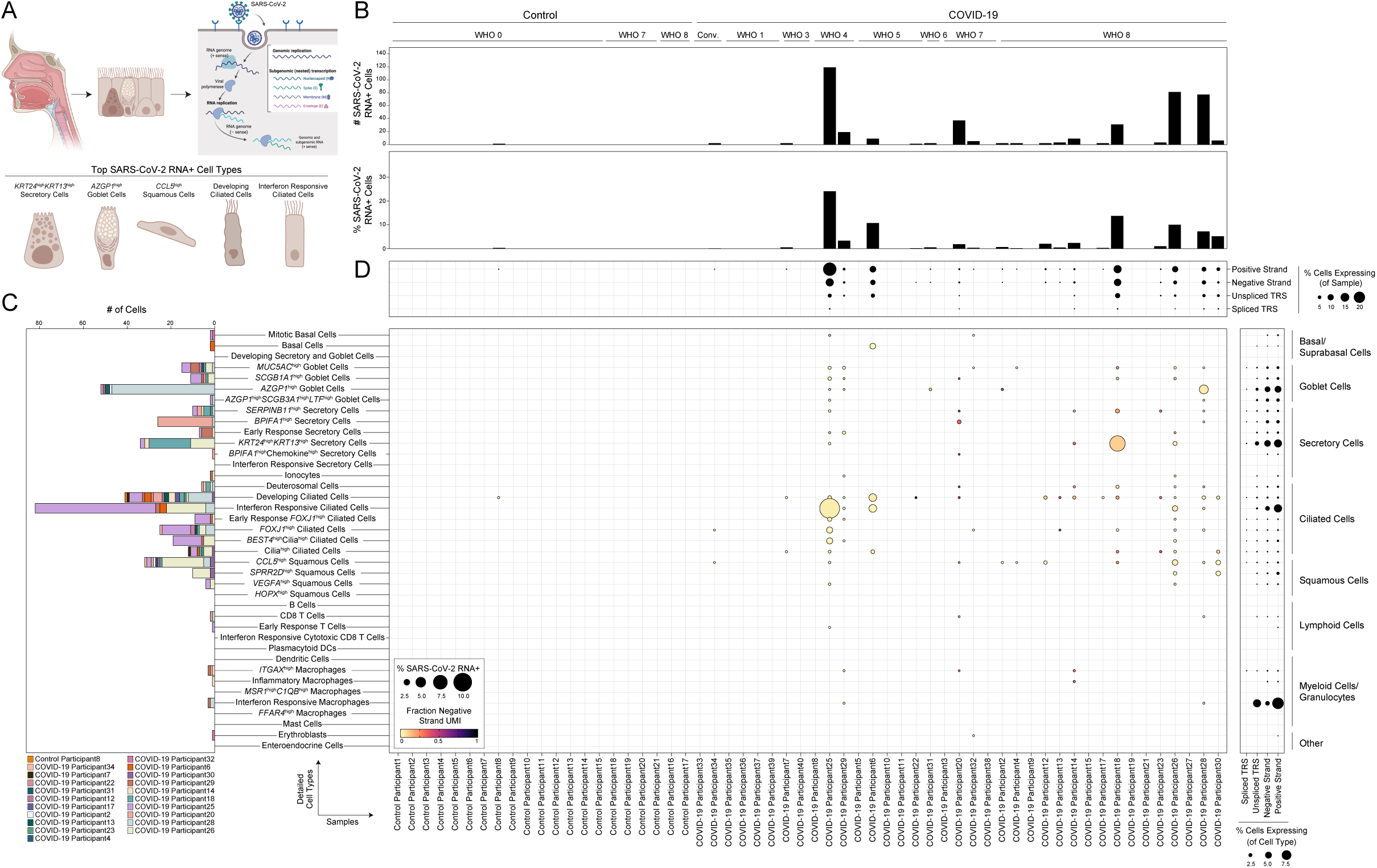
Cellular Targets of SARS-CoV-2 in the Nasopharynx. **A.** Summary schematic of top SARS-CoV-2 RNA+ cells. (created with BioRender). **B.** SARS-CoV-2 RNA+ cell number (top) and percent (bottom) per participant. Results following correction for ambient viral reads. **C.** Abundance of SARS-CoV-2 RNA+ cells by detailed cell type, bars colored by participant. Results following correction for ambient viral reads. **D.** Dot plot of SARS-CoV-2 RNA presence by sample (columns) and detailed cell types (rows). Dot size reflects fraction of a given participant and cell type containing SARS-CoV-2 RNA (following viral ambient correction). Dot color reflects fraction of aligned reads corresponding to the SARS-CoV-2 positive strand (yellow) vs. negative strand (black). Top dot plot across columns: alignment of viral reads by participant, separated by RNA species type. Right dot plot across rows: alignment of viral reads by detailed cell type, separated by RNA species type. *See also Figures S5, S6*

Finally, we compared the relative abundance of viral RNA *within* each cell type and found developing ciliated cells contain significantly higher SARS-CoV-2 RNA molecules per-cell, including positive strand, negative strand-aligning reads, and spliced TRS reads (**Supplementary Figure 6G**). Intriguingly, among ciliated cell subtypes, interferon responsive ciliated cells, despite representing one of the most frequent “targets” of viral infection, contain the lowest per-cell abundances of SARS-CoV-2 RNA, potentially reflecting the impact of elevated anti-viral factors curbing high levels of intracellular viral replication (**Supplementary Figure 6H**).

### Cell Intrinsic Responses to SARS-CoV-2 Infection

Above, we carefully mapped the specific cell types and states harboring SARS-CoV-2 RNA+ cells, identifying the subsets of epithelial cells that appear to actively support viral replication *in vivo* across distinct individuals (**Figure 5**). Further, we characterized robust and cell-type-specific host responses among cells from COVID-19 participants, ostensibly representing both the bystander cell response to local virus and an inflammatory microenvironment, as well as the intrinsic response to intracellular SARS-CoV-2 RNA (**Figure 3**). Here, by directly comparing single cells containing SARS-CoV-2 RNA to their matched bystanders, we aimed to map both the cell-intrinsic response to direct viral infection, as well as the host cell identities that may *potentiate* or *enable* SARS-CoV-2 tropism and replication.

To control for variability among different SARS-CoV-2 RNA+ cell types and individuals, we compared SARS-CoV-2 RNA+ cells to bystander cells of the same cell type and participant. Among cell types with at least 5 SARS-CoV-2 RNA+ cells, we observe robust and specific transcriptional changes compared to both matched bystander cells as well as cells from healthy individuals (**Figures 6A, 6B**). Notably, many of the genes previously identified as increased within all cells from COVID-19 participants, e.g., anti-viral factors *IFITM3, IFI44L*, are also upregulated among SARS-CoV-2 RNA+ cells compared to matched bystanders within multiple cell types. SARS-CoV-2 RNA+ cells from participants with mild/moderate COVID-19 show stronger induction of anti-viral and interferon responsive pathways compared to those from participants with severe COVID-19, despite equivalent abundances of cell-associated viral UMI (**Supplementary Figure 7A**). *EIF2AK2,* which encodes protein kinase R and drives host cell apoptosis following recognition of intracellular double-stranded RNA, is among the most reliably expressed and upregulated genes among SARS-CoV-2 RNA+ cells compared to matched bystanders across diverse cell types, suggesting rapid activation of this locus following intrinsic PAMP recognition of SARS-CoV-2 replication intermediates^97^. Therefore, direct sensing of intracellular viral products amplifies interferon-responsive and anti-viral gene upregulation, though these pathways are also elevated within bystander cells.

**Figure 6.**
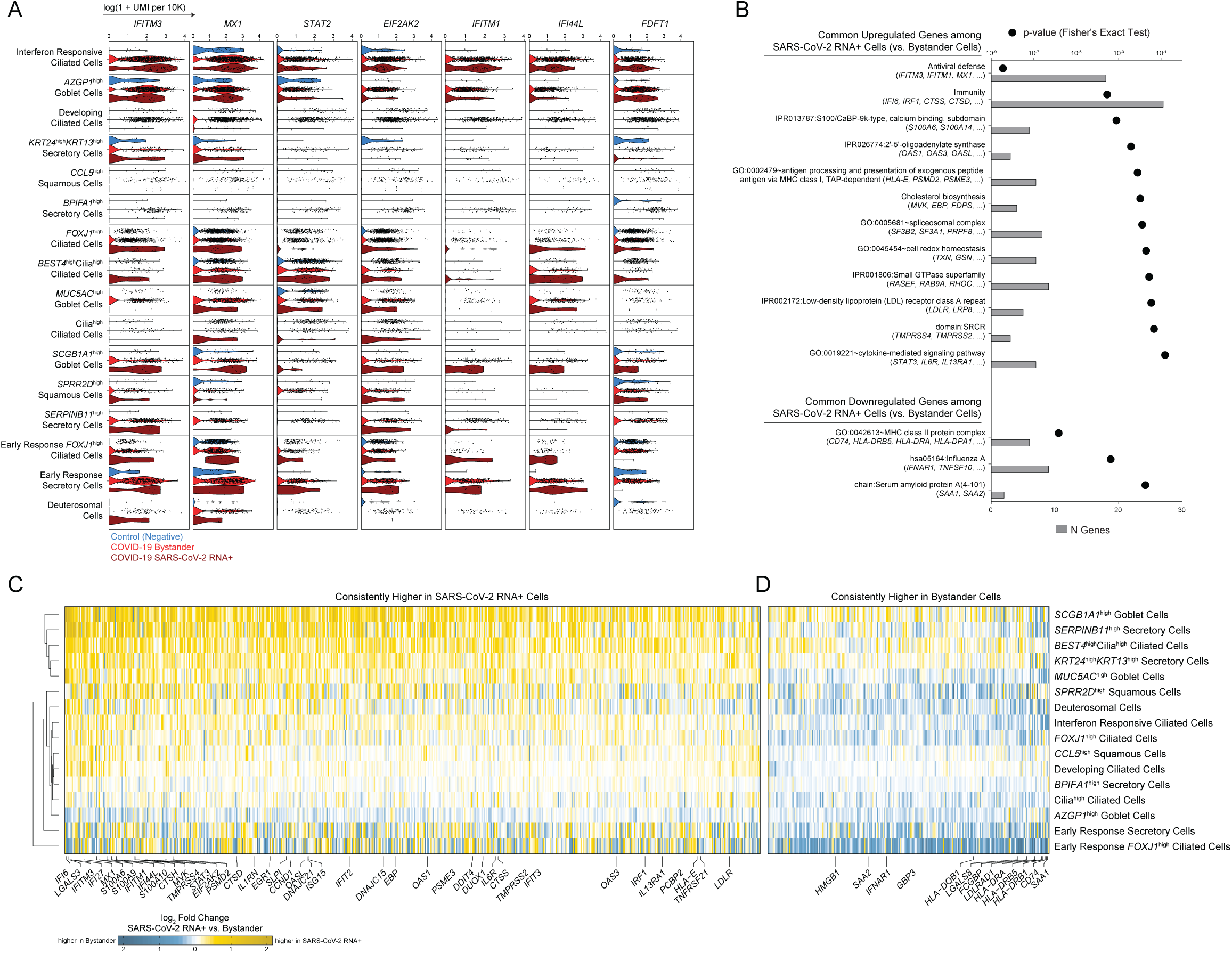
Intrinsic and Bystander Responses to SARS-CoV-2 Infection. **A.** Violin plots of selected genes upregulated in SARS-CoV-2 RNA+ cells in at least 3 individual cell type comparisons. Blue: cells from Control participants, Red: bystander cells from COVID-19 participants, Dark red: SARS-CoV-2 RNA+ cells. **B.** Enriched gene ontologies among genes consistently up- or down-regulated among SARS-CoV-2 RNA+ cells across cell types. **C.** Heatmap of genes consistently higher in SARS-CoV-2 RNA+ cells across multiple cell types. Colors represent log fold changes between SARS-CoV-2 RNA+ cells and bystander cells (SARS-CoV-2 RNA negative cells, from COVID-19 participants) by cell type. Restricted to cell types with at least 5 SARS-CoV-2 RNA+ cells. Yellow: upregulated among SARS-CoV-2 RNA+ cells, blue: upregulated among bystander cells. **D.** Heatmap of genes consistently higher in bystander cells across multiple cell types. *See also Figure S7, Table S5*

The majority of genes induced within SARS-CoV-2 RNA+ cells are shared *across* diverse cell types, suggesting a conserved anti-viral response, as well as common features that facilitate or restrict infection (**Figures 6B-6D**). SARS-CoV-2 RNA appears to robustly stimulate expression of genes involved in anti-viral sensing and defense (e.g., *MX1, IRF1, OAS1, OAS2)*, as well as genes involved in antigen presentation via MHC class I (**Figure 6C, Supplementary Table 5**). SARS-CoV-2 RNA+ cells expressed significantly higher abundances of multiple proteases involved in the cleavage of SARS-CoV-2 spike protein, a required step for viral entry (*TMPRSS4, TMPRSS2, CTSS, CTSD*). This suggests that within a given cell type, natural variations in the abundance of genes which support the viral life cycle may partially account for which cells are successfully targeted by the virus. Among the core anti-viral/interferon-responsive gene sets induced within SARS-CoV-2 RNA+ cells, we observed repeated and robust upregulation of *IFITM3* and *IFITM1*. Multiple studies have demonstrated that while these two interferon-inducible factors can disrupt viral release from endocytic compartments among a wide diversity of viral species, IFITMs can instead facilitate entry by human betacoronaviruses^95,98^. Therefore, enrichment of these factors within presumptive infected cells may reflect viral hijacking of a conserved host anti-viral responsive pathway. Genes involved in cholesterol and lipid biosynthesis are also upregulated among SARS-CoV-2 RNA+ cells, including *FDFT1*, *MVK, FDPS, ACAT2, HMGCS1*, all enzymes involved in the mevalonate synthesis pathway. In addition, SARS-CoV-2 RNA+ cells show increased abundance of low-density lipoprotein receptors *LDLR* and *LRP8* compared to matched bystanders. Intriguingly, various genes involved in cholesterol metabolism were recently identified as critical host factors for SARS-CoV-2 replication via CRISPR screens from multiple independent research groups^99,100^. Further, these groups found that direct inhibition of cholesterol biosynthesis decreased SARS-CoV-2 (as well as coronavirus strains 299E and OC43) replication within cell lines, and suggest S-mediated entry relies on host cholesterol. When we queried the full collections of presumptive replication factors identified by four published CRISPR screens^99–102^, we observed significant enrichment among SARS-CoV-2 RNA+ cells for RAB GTPases (e.g. *RAB9A, RHOC, RASEF*), vacuolar ATPase H+ pump subunits, as well as transcriptional modulators such as *SPEN, SLTM, CREBBP, SMAD4 and EGR1* (**Supplementary Figure 7B**).

Finally, we found multiple genes implicated in susceptibility and response to SARS-CoV-2 infection which have not been previously described, including S100/Calbindin genes such as *S100A6, S100A4,* and *S100A9*, which may directly play a role in leukocyte recruitment to infected cells. *IFNAR1* was substantially increased in many bystander cells compared to both cells from Control participants as well as matched SARS-CoV-2 RNA+ cells (**Figure 6D**). Blunting of interferon alpha signaling via downregulation of *IFNAR1* within SARS-CoV-2 RNA+ cells may partially explain high levels of viral replication compared to neighboring cells. Finally, bystander cells expressed significantly higher abundances of MHC-II molecules compared to SARS-CoV-2 RNA+ cells, including *HLA-DQB1, HLA-DRB1, HLA-DRB5, HLA-DRA,* and *CD74*.

## DISCUSSION

We present a comprehensive map of SARS-CoV-2 infection of the human nasopharynx using scRNA-seq. We hypothesize that the host response at the site of initial infection, the nasal mucosa, is an essential determinant of overall COVID-19 disease trajectory. By dissecting the nature of host-pathogen interactions at this primary viral target across the spectrum of disease outcomes, we can characterize both protective and pathogenic responses to SARS-CoV-2 infection. This enables us to begin to untangle the myriad factors that may restrict viral infection to the upper respiratory tract or support the development of severe lower respiratory tract disease.

First, we find that mature ciliated cells decline dramatically within the nasopharynx of COVID-19 samples, directly correlated with the tissue abundance of SARS-CoV-2 RNA at the time of sampling. Conversely, secretory cell populations expand among samples with high viral loads, which potentially represents a conserved response for epithelial re-population of lost mature ciliated cells through a recently-identified mechanism of secretory/goblet differentiation, using deuterosomal cells as intermediates^64,65^. Accordingly, deuterosomal cells and immature/developing ciliated cells are considerably expanded among COVID-19 samples, suggesting interdependence between each of these compartments in maintaining epithelial homeostasis during viral challenge. SARS-CoV-2 infection also induces dramatic increases in the diversity of epithelial cell types, both with respect to shifted compositional balance among major cell identities, and also via expansion of specialized secretory and goblet cell subsets, including a subset termed *KRT24*^high^*KRT13*^high^ secretory cells, which closely match the recently-identified KRT13+ “hillock” cell, previously associated with epithelial regions experiencing rapid cellular turnover and inflammation^65–67^. Other specialized subsets of secretory and goblet cells, such as early response secretory cells, *AZGP1*^high^ goblet cells, and *SCGB1A1*^high^ goblet cells, are expanded among COVID-19 participants. However, we observe expansion of these cells within discrete subsets of individuals, and they are not homogenous across the disease groups we sampled here. Indeed, understanding whether heterogeneous responses in the epithelial compartment *between* individuals with COVID-19 underscores differences in disease manifestations will require larger cohort studies, with a focus on longitudinal responses following initial infection. Further work is required to understand how the epithelial responses to SARS-CoV-2 infection within the nasal mucosa relates to epithelial responses during other upper respiratory viral infections and inflammatory states^84,103^.

Beyond cellular compositional changes during COVID-19, our study identifies marked variability in the induction of anti-viral gene expression that is associated with disease severity. We find robust upregulation of interferon stimulated genes among epithelial and immune cells isolated from individuals with mild/moderate COVID-19, and this is particularly evident in cells that contain SARS-CoV-2 RNA. Surprisingly, despite strong induction of anti-viral gene expression, we find little to no mRNA corresponding to type I or type III interferons amongst any recovered cell types. In a related study mapping the nasal epithelium during influenza infection, we and our colleagues observe extensive upregulation of *IFNA*, *IFNB1,* and *IFNL1-3* within ciliated cells and goblet cells, highlighting not only the capacity of superficial nasal epithelial cells to secrete local interferons during viral infection, but also the technical capacity of the scRNA-seq platform used in both studies to capture interferon mRNA^84^. The precise sources and signals which motivate a broad anti-viral response among mild COVID-19 cases in our study remain unknown – they may originate from immune cells contained deeper within the respiratory mucosa (therefore inaccessible through the superficial sampling used here), from sparse, highly transient interferon expression from superficial epithelial or immune cells, or may derive from direct PAMP/DAMP sensing or alternative inflammatory signals.

Remarkably, in comparison to individuals with mild/moderate disease, we find that anti-viral gene expression is profoundly blunted in cells isolated from individuals with severe disease, even in cells containing SARS-CoV-2 RNA. This effect is observed among diverse cell types, including those thought to represent direct targets of viral infection, such as ciliated cells and secretory cells, and also bystanders and co-resident immune cells. Notably, individuals with severe COVID-19 disease have equivalent or even elevated levels of nasal SARS-CoV-2 RNA at the time of sampling, and contained expanded inflammatory and type II-interferon responsive macrophages compared to mild/moderate cases. Indeed, published peripheral immune studies comparing mild and severe COVID-19 also observe diminished type I and type III interferon abundances in severe cases, and note restricted interferon stimulated gene expression among circulating immune cells^21–23^. Other human betacoronaviruses including MERS and SARS-CoV exhibit multiple strategies to avoid triggering pattern recognition receptor pathways, including degradation of host mRNA within infected cells^104,105^, sequestration of viral replication intermediates (e.g., double stranded RNA) from host sensors^106^, and direct inhibition of immune effector molecules^95,97,107^, thereby leading to diminished induction of anti-viral pathways and blunted autocrine and paracrine interferon signaling. Strategies to avoid innate immune recognition have now been extended to SARS-CoV-2 as well, indicating that avoiding host recognition is likely an essential aspect of viral success^108–110^. The close association we observe between disease severity and weak anti-viral gene expression among nasal epithelial cells is intriguing given recent observations of inborn defects in TLR3, IRF7, IRF9, and IFNAR1, or antibody-mediated neutralization of type I interferon responses within individuals who develop severe COVID-19^57–59^. Taken together, these findings strongly suggest that severe infection can arise in the setting of an intrinsic impairment of epithelial anti-viral immunity, and that timely induction of anti-viral responses is an essential aspect of successful viral control. We surmise that the combined effects of a viral strain with naturally poor interferon induction and intrinsic defects in immune or epithelial anti-viral responses within the nasal mucosa may predispose to severe disease via enhanced viral replication in the upper airway, eventually leading to immunopathology characteristic of severe COVID-19.

Critically, our work does not address the dynamics of nasal epithelial anti-viral responses during SARS-CoV-2 infection in individual patients, nor does it directly relate intrinsic mucosal responses in the nasopharynx to potential interferon or anti-viral responses in the lung or distal airways. Very recent work in this area profiling the earliest steps of host-virus interaction via nasopharyngeal transcriptional profiling and airway organoid models found that SARS-CoV-2 replicates with a doubling time of approximately 6 hours until interferon defenses begin to limit replication after ∼ 2 days^39^. However, different types of interferon may promote distinct host outcomes depending on when and where they are induced, as related work suggests type III interferons are present in the lungs, but not the nasopharynx, during SARS-CoV-2 infection, and may contribute to tissue damage late in disease course^111,112^. Further, while individuals in our cohort were not sampled during the asymptomatic phase, they were intentionally sampled as early within their disease course as possible and the majority have elevated viral levels within their nasopharynx, suggesting that a muted interferon response in severe patients can occur early even when viral loads are still matched with mild/moderate individuals. Further work will be required to directly link the nasopharyngeal profiles presented here to tissue and peripheral response during the hyper-inflammatory “late” stages of severe COVID-19. However, among individuals who develop severe COVID-19 in our cohort, we observe unique recruitment of highly inflammatory macrophages that represent the major tissue sources of proinflammatory cytokines including *IL1B*, *TNF*, *CXCL8*, *CCL2*, *CCL3* and *CXCL9/10/11* – of likely relation to the immune dysregulation characterized by elevation of the same factors in the periphery in late, severe disease and observe in lung tissue among those who succumbed to COVID-19^88^. In addition, we note specific upregulation of alarmins *S100A8/S100A9* (i.e., calprotectin) among epithelial cells in severe COVID-19 compared to mild and control counterparts, and even higher expression of *S100A9* within SARS-CoV-2 RNA+ cells from those same individuals. A recent study identified these as potential biomarkers of severe COVID-19, and proposed that these factors directly drive excessive inflammation and precede the massive cytokine release characteristic of late disease^113^. Our work suggests that severe COVID-19-specific expression of calprotectin may originate within the virally-infected nasal epithelia, and suggests that further work to understand the epithelial cell regulation of *S100A8/A9* gene expression may help clarify maladaptive responses to SARS-CoV-2 infection.

Finally, we provide a direct investigation into the host factors that enable or restrict SARS-CoV-2 replication within epithelial cells *in vivo*. Here, we recapitulate expected “hits” based on well-described host factors involved in viral replication – e.g., *TMPRSS2*, *TMPRSS4* enrichment among presumptive virally infected cells. We similarly observe expression of anti-viral genes which are globally enriched among cells from mild/moderate COVID-19 participants, with even higher expression among the viral RNA+ cells themselves. In accordance with previous studies into the nasal epithelial response to influenza infection^84^, we observe bystander epithelial cell upregulation of both MHC-I and MHC-II family genes; however, we find that SARS-CoV-2 RNA+ cells only express MHC-I, and poorly express MHC-II genes compared to matched bystanders. To our knowledge, downregulation of host cell pathways for antigen presentation by coronaviruses has not been previously described. A recent study found that CIITA and CD74 can intrinsically block entry of a range of viruses (including SARS-CoV-2) via endosomal sequestration, and therefore cells that upregulate these (and other) components of MHC-II machinery may naturally restrict viral entry^114^.

Together, our work demonstrates that many of the factors associated with the clinical trajectory following SARS-CoV-2 infection may stem from initial host-viral encounters in the nasopharyngeal epithelium. Further, it suggests that there may be a clinical window in which severe disease can be subverted by focusing preventative or therapeutic interventions early within the nasopharynx^115–118^, thereby bolstering anti-viral responses and curbing pathological inflammatory signaling prior to development of severe respiratory dysfunction or systemic disease.

## ACKNOWLEDGEMENTS

We thank the study participants and their families for enabling this research, the clinical support staff at UMMC for assistance in sample collection, and the members of the Shalek, Ordovas-Montanes, Horwitz, and Glover labs for thoughtful discussion and comments on the project. This project has been made possible in part by grant number 2020-216949 from the Chan Zuckerberg Initiative DAF, an advised fund of Silicon Valley Community Foundation to A.K.S. and J.O.M.. C.G.K.Z. was supported by T32GM007753 from the NIH. J.O.M is a New York Stem Cell Foundation – Robertson Investigator. J.O.M was supported by the Richard and Susan Smith Family Foundation, the AGA Research Foundation’s AGA-Takeda Pharmaceuticals Research Scholar Award in IBD – AGA2020-13-01, the Food Allergy Science Initiative, and The New York Stem Cell Foundation. B.H.H. was supported by DK122532-01A1, NIH; 12019PG-CD002, The Leona M. and Harry B. Helmsley Charitable Trust; SRA #54518, and Crohn’s and Colitis Foundation. P.L. was supported by 5T32AG000222 from the NIH. A.W.N. was supported by the Ludwig Center for Molecular Oncology at MIT. A.K.S. was supported by the Bill and Melinda Gates Foundation, Sloan Fellowship in Chemistry, the NIH (5U24AI118672), and the Ragon Institute of MGH, MIT and Harvard. We thank Joshua Gould, Katherine Siddle, Bo Li, Stephen Fleming, and the Broad Institute viral-ngs and Cumulus teams for assistance with computational pipelines.

## AUTHOR CONTRIBUTIONS

Conceptualization: J.O.-M., A.K.S., S.C.G., B.H.H.

Methodology: C.G.K.Z., V.N.M., A.W.N., Y.T., T.O.R., S.C.G., J.O.-M., A.K.S., B.H.H.

Formal analysis: C.G.K.Z., Y.T., A.H.O., P.L.

Investigation: C.G.K.Z., V.N.M., A.W.N., Y.T., J.D.B., M.G., R.S.D., H.B.W., M.Sl., A.H.O., M.H., H.L.,

Resources: S.C.G., A.H.O., M.Sl., H.L., A.P., C.B.S., M.W.B., M.Se., M.H., G.E.A.

Data Curation: T.O.R., Y.T., M.Sl. A.H.O., H.L., H.B.W., T.C., C.G.K.Z.

Writing – Original Draft: C.G.K.Z., J.O.-M., A.K.S., S.C.G., B.H.H.

Writing – Review & Editing: C.G.K.Z., V.N.M., A.H.O., A.W.N., Y.T., J.D.B., P.L., M.Sl., H.L., H.B.W., M.G.,

R.S.D., T.C., A.P., C.B.S., M.W.B., Y.P., M.H., G.E.A., M.Se., T.O.R., A.K.S., S.C.G., B.H.H., J.O.-M.

Visualization: C.G.K.Z., Y.T.

Supervision: J.O.-M., A.K.S., S.C.G., B.H.H., T.O.R.

Project Administration: Y.P., T.O.R.

Funding Acquisition: J.O.-M., A.K.S., S.C.G., B.H.H.

## METHODS

### Study Participants and Design

Eligible participants were recruited from outpatient clinics, medical surgical units, intensive care units (ICU), or endoscopy units at the University of Mississippi Medical Center (UMMC) between April 2020 and September 2020. The UMMC Institutional Review Board approved the study under IRB#2020-0065. All participants or their legally authorized representative provided written informed consent. Participants were eligible for inclusion in the COVID-19 group if they were at least 18 years old, had a positive nasopharyngeal swab for SARS-CoV-2 by PCR, had COVID-19 related symptoms including fever, chills, cough, shortness of breath, and sore throat, and weighed more than 110 lb. Participants were eligible for the Control group if they were at least 18 years old, had a current negative SARS-CoV-2 test (PCR or rapid antigen test), and weighed more than 110 lb. Participants were considered “Convalescent” if they met the criteria of the Control group, however had previously tested SARS-CoV-2 PCR positive and diagnosed with COVID-19, and their symptoms had subsided for at least 40 days. Convalescent samples were treated as an independent group, and excluded from comparisons between “Control” and “COVID-19” groups. Exclusion criteria for the cohort included a history of blood transfusion within 4 weeks and subjects who could not be assigned a definitive COVID-19 diagnosis from nucleic acid testing. 35 individuals with COVID-19 were included, both male (n = 19) and female (n = 16). For the Control group, 21 participants were included – 11 identified as male, 10 as female. The median age of COVID-19 participants was 55 years old; the median age of Control participants was 62 years old. Among hospitalized COVID-19 participants, the median day NP swabs were collected was hospital day 2 (inter-quartile range: 1, range 1-9). COVID-19 participants were classified according to the 8-level ordinal scale proposed by the WHO representing their peak clinical severity and level of respiratory support required^60^. See **Table 1** and **Supplementary Figure 1A**.

### Sample Collection and Biobanking

Nasopharyngeal samples were collected by a trained healthcare provider using FLOQSwabs (Copan flocked swabs) following the manufacturer’s instructions. Collectors would don personal protective equipment (PPE), including a gown, non-sterile gloves, a protective N95 mask, a bouffant, and a face shield. The patient’s head was tilted back slightly, and the swab inserted along the nasal septum, above the floor of the nasal passage to the nasopharynx until slight resistance was felt. The swab was then left in place for several seconds to absorb secretions and slowly removed while rotating swab. The swab was then placed into a cryogenic vial with 900 µL of heat inactivated fetal bovine serum (FBS) and 100 µL of dimethyl sulfoxide (DMSO). Vials were placed into a Mr. Frosty Freezing Container (Thermo Fisher Scientific) for optimal cell preservation. A Mr. Frosty containing the vials was placed in a cooler with dry ice for transportation from patient areas to the laboratory for processing. Once in the laboratory, the Mr. Frosty was placed into a −80°C freezer overnight, and on the next day, the vials were moved to liquid nitrogen storage containers.

### Dissociation and Collection of Viable Single Cells from Nasopharyngeal Swabs

Swabs in freezing media (90% FBS/10% DMSO) were stored in liquid nitrogen until immediately prior to dissociation. A detailed sample protocol can be found here: https://protocols.io/view/human-nasopharyngeal-swab-processing-for-viable-si-bjhkkj4w.html119. This approach (**Supplementary Figure 1C**) ensures that all cells and cellular material from the nasal swab (whether directly attached to the nasal swab, or released during the washing and digestion process), are exposed first to DTT for 15 minutes, followed by an Accutase digestion for 30 minutes. Briefly, nasal swabs in freezing media were thawed, and each swab was rinsed in RPMI before incubation in 1 mL RPMI/10 mM DTT (Sigma) for 15 minutes at 37°C with agitation. Next, the nasal swab was incubated in 1 mL Accutase (Sigma) for 30 minutes at 37°C with agitation. The 1 mL RPMI/10 mM DTT from the nasal swab incubation was centrifuged at 400 g for 5 minutes at 4°C to pellet cells, the supernatant was discarded, and the cell pellet was resuspended in 1 mL Accutase and incubated for 30 minutes at 37°C with agitation. The original cryovial containing the freezing media and the original swab washings were combined and centrifuged at 400 g for 5 minutes at 4°C. The cell pellet was then resuspended in RPMI/10 mM DTT, and incubated for 15 minutes at 37°C with agitation, centrifuged as above, the supernatant was aspirated, and the cell pellet was resuspended in 1 mL Accutase, and incubated for 30 minutes at 37°C with agitation. All cells were combined following Accutase digestion and filtered using a 70 µm nylon strainer. The filter and swab were washed with RPMI/10% FBS/4 mM EDTA, and all washings combined. Dissociated, filtered cells were centrifuged at 400 g for 10 minutes at 4°C, and resuspended in 200 µL RPMI/10% FBS for counting. Cells were diluted to 20,000 cells in 200 µL for scRNA-seq. For the majority of swabs, fewer than 20,000 cells total were recovered. In these instances, all cells were input into scRNA-seq.

### Flow Cytometry of Cells Isolated from Nasopharyngeal Swabs

Single-cell suspensions were isolated from nasopharyngeal swabs of healthy donors, as described above. Cells were first stained with Fixable Aqua Dead Cell Stain (Thermo Fisher Scientific) for 15 mins to assess viability. Cells were washed with staining buffer (PBS/2% FBS), and then treated with Human TruStain FcX (Fc receptor blocking solution, Cat. No. 422302, BioLegend) for 5 mins. Cells were then stained with a surface marker antibody cocktail on ice for 15 mins, which contained PerCP-Cy5.5-conjugated anti-human CD45 (clone: HI30, BioLegend), Brilliant Violet 711-conjugated anti-human CD3 (clone: SK7, BioLegend), APC-Cy7-conjugated anti-human CD8 (clone: SK1, BioLegend), PE-conjugated anti-human CD4 (clone: RPA-T4, BioLegend), Brilliant Violet 786-conjugated anti-human CD326 (clone: 9C4, BioLegend), PE-Cy5-conjugated anti-human CD19 (clone: HIB19, BioLegend), PE-Cy7-conjugated anti-human CD66b (clone: G10F5, BioLegend), Brilliant Violet 650-conjugated anti-human CD11c (clone: Bu15, BioLegend), FITC-conjugated anti-human CD14 (clone: M5E2, BioLegend), and Brilliant Violet 421-conjugated anti-human CD56 (clone: 5.1H11, BioLegend). Finally, cells were fixed using BD Fixation Buffer (Cat. No. 554655, BD Biosciences). AbC Total Antibody Compensation Beads (Cat. No. A10497, Life Technologies) and ArC Amine Reactive Compensation Beads (Cat. No. A10346, Life Technologies) were used for compensation. Data were acquired on an LSRFortessa flow cytometer (BD Biosciences) using BD FACSDiva software, and analyzed by FlowJo software (Version 10.7.1, Tree Star Inc.).

### Single Cell RNA-Sequencing

Seq-Well S^3^ was run as previously described^61,62,120^. Briefly, a maximum of 20,000 single cells were deposited onto Seq-Well arrays preloaded with a single barcoded mRNA capture bead (ChemGenes) per well^121^. cells were allowed to settle by gravity into wells for 10 minutes, after which the arrays were washed with PBS and RPMI, and sealed with a semi-permeable membrane for 30 minutes, and incubated in lysis buffer (5 M guanidinium thiocyanate/1 mM EDTA/1% BME/0.5% sarkosyl) for 20 minutes. Arrays were then incubated in a hybridization buffer (2M NaCl/8% v/v PEG8000) for 40 minutes, and then the beads were removed from the arrays and collected in 1.5 mL tubes in wash buffer (2M NaCl/3 mM MgCl_2_/20 mM Tris-HCl/8% v/v PEG8000). Beads were resuspended in a reverse transcription master mix, and reverse transcription, exonuclease digestion, second-strand synthesis, and whole transcriptome amplification were carried out as previously described. Libraries were generated using Illumina Nextera XT Library Prep Kits and sequenced on NextSeq 500/550 High Output 75 cycle v2.5 kits to an average depth of 180 million aligned reads per array: read 1: 21 (cell barcode, UMI), read 2: 50 (digital gene expression), index 1: 8 (N700 barcode).

### Data Preprocessing and Quality Control

Pooled libraries were demultiplexed using bcl2fastq (v2.17.1.14) with default settings (mask_short_adapter_reads 10, minimum_trimmed_read_length 10, implemented using Cumulus, snapshot 4, https://cumulus.readthedocs.io/en/stable/bcl2fastq.html)122. Libraries were aligned using STAR within the Drop-Seq Computational Protocol (https://github.com/broadinstitute/Drop-seq) and implemented on Cumulus (https://cumulus.readthedocs.io/en/latest/drop_seq.html, snapshot 9, default parameters)^121^. A custom reference was created by combining human GRCh38 (from CellRanger version 3.0.0, Ensembl 93) and SARS-CoV-2 RNA genomes. The SARS-CoV-2 viral sequence and GTF are as described in Kim et al. 2020 (https://github.com/hyeshik/sars-cov-2-transcriptome, BetaCov/South Korea/KCDC03/2020 based on NC_045512.2)^87^. The GTF includes all CDS regions (as of this annotation of the transcriptome, the CDS regions completely cover the RNA genome without overlapping segments), and regions were added to describe the 5’ UTR (“SARSCoV2_5prime”), the 3’ UTR (“SARSCoV2_3prime”), and reads aligning to anywhere within the Negative Strand (“SARSCoV2_NegStrand”). Trailing A’s at the 3’ end of the virus were excluded from the SARS-CoV-2 FASTA, as these were found to drive spurious viral alignment in pre-COVID19 samples. Finally, additional small sequences were appended to the FASTA and GTF that differentiate reads that align to the 70-nucleotide region around the viral TRS sequence – either across the intact, unspliced genomic sequences (e.g. named “SARSCoV2_Unspliced_S” or “SARSCoV2_Unspliced_Leader”) or various spliced RNA species (e.g. “SARSCoV2_Spliced_Leader_TRS_S”), see schematics in **Figures 4F, 4G, Supplementary Figures 6A**. Alignment references were tested against a diverse set of pre-COVID-19 samples and *in vitro* SARS-CoV-2 infected human bronchial epithelial cultures^37^ to confirm specificity of viral aligning reads. Aligned cell-by-gene matrices were merged across all study participants, and cells were filtered to eliminate barcodes with fewer than 200 UMI, 150 unique genes, and greater than 50% mitochondrial reads. Swabs from 58 individuals are included in the study. Three additional swabs were thawed and processed but contained no high-quality cell barcodes after sequencing (**NB**: these samples contained < 5,000 viable cells prior to Seq-Well array loading).This resulted in a final dataset of 32,871 genes and 32,588 cells across 58 study participants (35 SARS-CoV-2+, 23 SARS-CoV-2-). Preprocessing, alignment, and data filtering was applied equivalently to samples from the fresh vs. frozen cohort. For analysis of RNA velocity, we also recovered both exonic and intronic alignment information using DropEst (Cumulus (https://cumulus.readthedocs.io/en/latest/drop_seq.html, snapshot 9, dropest_velocyto true, run_dropest true)^123^.

### Cell Clustering and Annotation

Dimensionality reduction, cell clustering, and differential gene analysis were all achieved using the Seurat (v3.1.5) package in R programming language (v3.0.2)^124^. Dimensionality reduction was carried out by running principal components analysis (PCA) over the 3,483 most variable genes with dispersion > 0.8 (tested over a range of dispersion > 0.7 to dispersion > 1.2; dispersion > 0.8 was determined as optimal based on number of variable genes, and general stability of clustering results across these cutoffs was confirmed). Only variable genes from human transcripts were considered for dimensionality reduction and clustering. Using the Jackstraw function within Seurat, we selected the first 36 principal components that described the majority of variance within the dataset, and used these for defining a nearest neighbor graph and Uniform Manifold Approximation and Projection (UMAP) plot. Cells were clustered using Louvain clustering, and the resolution parameter was chosen by maximizing the average silhouette score across all clusters. Differentially expressed genes between each cluster and all other cells were calculated using a likelihood ratio test, implemented with Seurat’s FindAllMarkers function, test.use set to “bimod”^125^. Clusters were merged if they failed to contain > 25 significantly differentially expressed genes (FDR < 0.001). We proceeded iteratively through each cluster and subcluster until “terminal” cell subsets/cell states were identified – we defined “terminal” cell states when PCA and Louvain clustering did not confidently identify additional sub-states, as measured by abundance of differentially expressed genes between potential clusters (often > 25 cluster-specific marker genes with FDR < 0.001). Among two samples, we recovered erythroblast-like cells, defined by expression of hemoglobin subunits including *HBB and HBA2* (these were from swabs noted to be slightly red-tinged on day of processing). For visualization in **Figure 2**, we pooled all cells determined to be of epithelial origin from coarse-grained annotation (ciliated cells, secretory cells, goblet cells, basal cells, mitotic basal cells, developing secretory and goblet cells, developing ciliated cells, squamous cells, deuterosomal cells, and ionocytes) and used the methods for dimensionality reduction as above (dispersion cutoff > 1, 30 principal components). We applied similar approaches for immune cell types (**Supplementary Figure 3**), including iterative subclustering to resolve and annotate all constituent cells types and subtypes. Gene module scores were calculated using the AddModuleScore function within Seurat.

We annotated epithelial subtypes according to the following groups and representative markers: goblet cells were split into 4 distinct sets: *MUC5AC*^high^ goblet cells, which lacked additional specialized markers beyond classic goblet cell identifiers, *SCGB1A1*^high^ goblet cells, *AZGP1*^high^ goblet cells, and *AZGP1*^high^*SCGB3A1*^high^ *LTF*^high^ goblet cells. Secretory cells were divided into 6 distinct detailed subtypes: *SERPINB11*^high^ secretory cells (which, similar to *MUC5AC*^high^ goblet cells, represented a more “generic” or un-differentiated secretory cell phenotype), *BPIFA1*^high^ secretory cells, early response secretory cells (which expressed genes such as *JUN, EGR1, FOS, NR4A1*), *KRT24*^high^*KRT13*^high^ secretory cells, *BPIFA1*^high^chemokine^high^ secretory cells (chemokines include *CXCL8, CXCL2, CXCL1,* and *CXCL3),* and interferon responsive secretory cells (defined by higher expression of broad anti-viral genes including *IFITM3, IFI6,* and *MX1*). Subsets of squamous cells were also found – detailed squamous cell subtypes include *CCL5*^high^ squamous cells, *VEGFA*^high^ squamous cells (which express multiple vascular endothelial genes including *VEGFA and VWF*), *SPRR2D*^high^ squamous cells (which, in addition to *SPRR2D*, express the highest abundances of multiple SPRR-genes including *SPRR2A, SPRR1B, SPRR2E,* and *SPRR3*), and *HOPX*^high^ squamous cells. Finally, ciliated cells could be further divided into 5 distinct subtypes: interferon responsive ciliated cells (expressing anti-viral genes similar to other “interferon responsive” subsets, such as *IFIT1, IFIT3, IFI6*), *FOXJ1*^high^ ciliated cells, early response *FOXJ1*^high^ ciliated cells (which, in addition to high *FOXJ1*, also express higher abundances of genes such as *JUN*, *EGR1*, *FOS* than other ciliated cell subtypes), cilia^high^ ciliated cells (which broadly express the highest abundances of structural cilia genes, such as *DLEC1* and *CFAP100),* and *BEST4*^high^cilia^high^ ciliated cells (in addition to cilia components, also express the ion channel *BEST4*).

### RNA Velocity and Pseudotemporal Ordering of Epithelial Cells

RNA velocity was modeled using the scVelo package, version 0.2.3^77,78^. Briefly, RNA velocity analysis leverages the dynamic relationships between expression of unspliced (intron-containing) and spliced (exonic) RNA across thousands of variable genes, enabling 1) estimation of the directionality of transitions between distinct cells and cell types, and 2) identification of putative driver genes behind these transitions. Using cluster annotations previously assigned from iterative clustering in Seurat, cells from epithelial cell types were pre-processed according to the scVelo pipeline: genes were normalized using default parameters (pp.filter_and_normalize), principal components and nearest neighbors in PCA space were calculated (using defaults of 30 PCs, 30 nearest neighbors), and the first and second order moments of nearest neighbors were computed, which are used as inputs into velocity estimates (pp.moments). RNA velocity was estimated using the scVelo tool tl.recover_dynamics with default input parameters, which maps the full splicing kinetics for all genes and tl.velocity, with mode=’dynamical’. Top velocity transition “driver” genes were identified by high “fit_likelihood” parameters from the dynamical model, and are used for visualization in **Supplementary Figure 2C**. The same approaches were used for modeling RNA velocity among only basal, secretory, and goblet cells (**Figures 2F-2I**), only ciliated cells (**Figures 2K-2N**), and only COVID-19 or only Control cells (**Figures 2P, 2Q**). For RNA velocity analysis of ciliated cells or basal, secretory and goblet cells, the velocity pseudotime was calculated using the tl.velocity_pseudotime function with default settings.

### Metatranscriptomic Classification of Reads from Single-Cell RNA-Seq

To identify co-detected microbial taxa present in the cell-associated or ambient RNA of nasopharyngeal swabs, we used the Kraken2 software implemented using the Broad Institute viral-ngs pipelines on Terra (https://github.com/broadinstitute/viral-pipelines/tree/master)86. A previously-published reference database included human, archaea, bacteria, plasmid, viral, fungi, and protozoa species and was constructed on May 5, 2020, therefore included sequences belonging to the novel SARS-CoV-2 virus^85^. Inputs to Kraken2 were: kraken2_db_tgz = “gs://pathogen-public-dbs/v1/kraken2-broad-20200505.tar.zst”, krona_taxonomy_db_kraken2_tgz = “gs://pathogen-public-dbs/v1/krona.taxonomy-20200505.tab.zst”, ncbi_taxdump_tgz = “gs://pathogen-public-dbs/v1/taxdump-20200505.tar.gz”, trim_clip_db = “gs://pathogen-public-dbs/v0/contaminants.clip_db.fasta” and spikein_db = “gs://pathogen-public-dbs/v0/ERCC_96_nopolyA.fasta”. Viral species with fewer than 5 reads were considered spurious and excluded.

### Correction for Ambient Viral RNA

Data from high-throughput scRNA-seq platforms frequently experience low-levels of non-specific RNA assigned to cell barcodes that does not represent true cell-derived transcriptomic material, but rather contamination from the ambient pool of RNA. To safeguard against spurious assignment of SARS-CoV-2 RNA to cells without true intracellular viral material, i.e., viral RNA non-specifically picked up from the microenvironment as a component of ambient RNA contamination, we employed the following corrections and statistical tests to control for ambient viral RNA and enable confident assignments for SARS-CoV-2 RNA+ cells. Similar to approaches previously described, we tested whether the abundance of viral RNA within a given single cell was significantly higher than expected by chance given the estimate of ambient RNA contaminating that cell, as well as the proportion of viral RNA of the total ambient RNA pool^84,88,92,93^. First, this required modeling and estimating the ambient RNA fraction associated with each individual swab. Here, we employed CellBender (https://github.com/broadinstitute/CellBender), a software package built to learn the ambient RNA profile per sample and provide an ambient RNA-corrected output^93^. Input UMI count matrices contained the top 10,000 cell barcodes, therefore including at least 70% cell barcodes sampling the ambient RNA and low-complexity cell barcodes. CellBender’s remove-background function was run with default parameters and --fpr 0.01 --expected-cells 500 --low-count-threshold 5. Using the corrected output from each sample’s count matrix following CellBender, we calculated the proportion of ambient contamination per high-quality cell by comparing to the single-cell’s transcriptome pre-correction, and summed all UMI from background cell barcodes to recover an estimate of the total ambient pool. Next, we tested whether the abundance of viral RNA in a given single cell was significantly above the null abundance given the ambient RNA characteristics using an exact binomial test (implemented in R: binom.test):

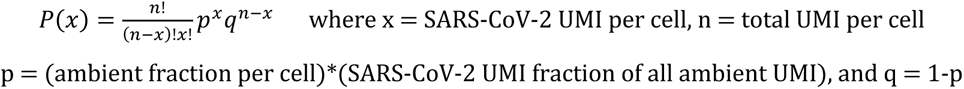

P-values were FDR-corrected within sample, and cells whose SARS-CoV-2 UMI abundance with FDR < 0.01 were considered “SARS-CoV-2 RNA+”.

### Differential Expression by Group, Cell Type, or Viral RNA Status

To compare gene expression between cells from distinct disease groups we employed a likelihood ratio test assuming a negative binomial distribution. Cells from each cell type belonging to either COVID-19 WHO 1-5 (mild/moderate), COVID-19 WHO 6-8 (severe), or Control WHO 0 were compared in a pairwise manner, implemented using the Seurat FindAllMarkers function (test.use = “negbinom”). We considered genes as differentially expressed with an FDR-adjusted p value < 0.001 and log fold change > 0.25. To compare gene expression between SARS-CoV-2 RNA+ cells and bystander cells (from COVID-19 participants, but without intracellular viral RNA) we used a negative binomial generalized linear model implemented using DESeq2^126^. Here, we employed the following criteria for SARS-CoV-2 RNA+ vs. bystander testing: 1) we only tested cell types containing at least 15 SARS-CoV-2 RNA+ cells, 2) for each cell type, we restricted our bystander cells to the same participants as the SARS-CoV-2 RNA+ cells, 3) in comparisons where bystander cells were substantially more numerous than SARS-CoV-2 RNA+ cells, we randomly sub-sampled the bystander cells to at most 4x the number of SARS-CoV-2 RNA+ cells, and 4) we ensured that the sampled bystander cells for comparison matched the cell quality distribution of the SARS-CoV-2 RNA+ cells, based on binned deciles of UMI/cell. DESeq2 was run with default parameters and test = “Wald”. Gene ontology analysis was run using the Database for Annotation, Visualization, and Integrated Discovery (DAVID)^127^. Gene set enrichment analysis (GSEA) was completed using the R package fgsea over genes ranked by average log foldchange expression between each group, including all genes with an average expression > 0.5 UMI within each respective cell type^128^. Gene lists corresponding to “Shared IFN Response”, “Type I IFN Specific Response” and “Type II IFN Specific Response” are derived from previously-published population RNA-seq data from nasal epithelial basal cells treated *in vitro* with 0.1 ng/mL – 10 ng/mL IFNα or IFNγ for 12 hours^29^. Module scores were calculated using the Seurat function AddModuleScore with default inputs.

### Statistical Testing

All statistical tests were implemented either in R (v4.0.2) or Prism (v6) software^129^. Comparisons between cell type proportions by disease group were tested using a Kruskal-Wallis test with FDR correction across all cell types, implemented in R using the kruskal.test, and p.adjust functions. Post-tests for between-group pairwise comparisons used Dunn’s test. Spearman correlation was used where appropriate, implemented using the cor.test function in R. All testing for differential expression was implemented in R using either Seurat, scVelo, or DESeq2, and all results were FDR-corrected as noted in specific **Methods** sections. P-values, n, and all summary statistics are provided either in the results section, figure legends, figure panels, or supplementary tables.

### Data and Code Availability

Prism (v6), R (v4.0.2) packages ggplot2 (v3.3.2^130^), Seurat (v3.2.2^131^), ComplexHeatmap (v2.7.3^132^), Circlize (0.4.11^133^), fgsea (v.1.16.0^128^), DESeq2 (v1.30.0^126^), and Python (v3.8.3) package scVelo (v0.3.0^77^) were used for visualization. All raw, normalized, and annotated data is available for download and visualization via the Single Cell Portal: https://singlecell.broadinstitute.org/single_cell/study/SCP1289/. Data was also deposited in a single-cell data resource for COVID-19 studies: https://www.covid19cellatlas.org134. Custom reference FASTA and GTF for SARS-CoV-2 is available for download: https://github.com/ShalekLab/SARSCoV2-genome-reference.

## REFERENCES

1. Pan, Y., Zhang, D., Yang, P., Poon, L. L. M. & Wang, Q. Viral load of SARS-CoV-2 in clinical samples. The Lancet Infectious Diseases (2020) doi:10.1016/S1473-3099(20)30113-4.

2. Sanche, S. et al. RESEARCH High Contagiousness and Rapid Spread of Severe Acute Respiratory Syndrome Coronavirus 2. Emerg. Infect. Dis. (2020) doi:10.3201/eid2607.200282.

3. Meyerowitz, E. A., Richterman, A., Gandhi, R. T. & Sax, P. E. Transmission of SARS-CoV-2: A Review of Viral, Host, and Environmental Factors. Ann. Intern. Med. (2021) doi:10.7326/m20-5008.

4. Fears, A. C. et al. Persistence of Severe Acute Respiratory Syndrome Coronavirus 2 in Aerosol Suspensions. Emerg. Infect. Dis. (2020) doi:10.3201/eid2609.201806.

5. Arons, M. M. et al. Presymptomatic SARS-CoV-2 Infections and Transmission in a Skilled Nursing Facility. N. Engl. J. Med. (2020) doi:10.1056/nejmoa2008457.

6. Wang, Y. et al. Clinical outcome of 55 asymptomatic cases at the time of hospital admission infected with SARS-Coronavirus-2 in Shenzhen, China. J. Infect. Dis. (2020) doi:10.1093/infdis/jiaa119.

7. Sakurai, A. et al. Natural History of Asymptomatic SARS-CoV-2 Infection. N. Engl. J. Med. (2020) doi:10.1056/nejmc2013020.

8. Guan, W. et al. Clinical Characteristics of Coronavirus Disease 2019 in China. N. Engl. J. Med. (2020) doi:10.1056/nejmoa2002032.

9. Huang, C. et al. Clinical features of patients infected with 2019 novel coronavirus in Wuhan, China. Lancet (2020) doi:10.1016/S0140-6736(20)30183-5.

10. Chan, J. F. W. et al. Genomic characterization of the 2019 novel human-pathogenic coronavirus isolated from a patient with atypical pneumonia after visiting Wuhan. Emerg. Microbes Infect. (2020) doi:10.1080/22221751.2020.1719902.

11. Zhou, P. et al. A pneumonia outbreak associated with a new coronavirus of probable bat origin. Nature (2020) doi:10.1038/s41586-020-2012-7.

12. Wu, F. et al. A new coronavirus associated with human respiratory disease in China. Nature (2020) doi:10.1038/s41586-020-2008-3.

13. Frieman, M. & Baric, R. Mechanisms of Severe Acute Respiratory Syndrome Pathogenesis and Innate Immunomodulation. Microbiol. Mol. Biol. Rev. (2008) doi:10.1128/mmbr.00015-08.

14. Harrison, A. G., Lin, T. & Wang, P. Mechanisms of SARS-CoV-2 Transmission and Pathogenesis. Trends in Immunology (2020) doi:10.1016/j.it.2020.10.004.

15. Borczuk, A. C. et al. COVID-19 pulmonary pathology: a multi-institutional autopsy cohort from Italy and New York City. Mod. Pathol. (2020) doi:10.1038/s41379-020-00661-1.

16. Ackermann, M. et al. Pulmonary Vascular Endothelialitis, Thrombosis, and Angiogenesis in Covid-19. N. Engl. J. Med. (2020) doi:10.1056/nejmoa2015432.

17. Lucas, C. et al. Longitudinal analyses reveal immunological misfiring in severe COVID-19. Nature (2020) doi:10.1038/s41586-020-2588-y.

18. Mathew, D. et al. Deep immune profiling of COVID-19 patients reveals distinct immunotypes with therapeutic implications. Science (80-. ). (2020) doi:10.1126/SCIENCE.ABC8511.

19. Schulte-Schrepping, J. et al. Severe COVID-19 Is Marked by a Dysregulated Myeloid Cell Compartment. Cell (2020) doi:10.1016/j.cell.2020.08.001.

20. Su, Y. et al. Multi-Omics Resolves a Sharp Disease-State Shift between Mild and Moderate COVID-19. Cell (2020) doi:10.1016/j.cell.2020.10.037.

21. Galani, I. E. et al. Untuned antiviral immunity in COVID-19 revealed by temporal type I/III interferon patterns and flu comparison. Nat. Immunol. (2021) doi:10.1038/s41590-020-00840-x.

22. Hadjadj, J. et al. Impaired type I interferon activity and inflammatory responses in severe COVID-19 patients. Science (80-. ). (2020) doi:10.1126/science.abc6027.

23. Stephenson, E. et al. The cellular immune response to COVID-19 deciphered by single cell multi-omics across three UK centres. medRxiv (2021).

24. Wilk, A. J. et al. A single-cell atlas of the peripheral immune response in patients with severe COVID-19. Nat. Med. (2020) doi:10.1038/s41591-020-0944-y.

25. Kusnadi, A. et al. Severely ill COVID-19 patients display impaired exhaustion features in SARS-CoV-2-reactive CD8 + T cells. Sci. Immunol. (2021) doi:10.1126/sciimmunol.abe4782.

26. Szabo, P. A. et al. Analysis of respiratory and systemic immune responses in COVID-19 reveals mechanisms of disease pathogenesis. medRxiv (2020) doi:10.1101/2020.10.15.20208041.

27. Ren, X. et al. COVID-19 immune features revealed by a large-scale single cell transcriptome atlas. Cell (2021).

28. Sungnak, W. et al. SARS-CoV-2 entry factors are highly expressed in nasal epithelial cells together with innate immune genes. Nat. Med. (2020) doi:10.1038/s41591-020-0868-6.

29. Ziegler, C. G. K. et al. SARS-CoV-2 Receptor ACE2 Is an Interferon-Stimulated Gene in Human Airway Epithelial Cells and Is Detected in Specific Cell Subsets across Tissues. Cell (2020) doi:10.1016/j.cell.2020.04.035.

30. Huang, N. et al. Integrated Single-Cell Atlases Reveal an Oral SARS-CoV-2 Infection and Transmission Axis. medRxiv (2020).

31. Muus, C. et al. Integrated analyses of single-cell atlases reveal age, gender, and smoking status associations with cell type-specific expression of mediators of SARS-CoV-2 viral entry and highlights inflammatory programs in putative target cells. bioRxiv 2020.04.19.049254 (2020) doi:10.1101/2020.04.19.049254.

32. Lukassen, S. et al. SARS-CoV-2 receptor ACE2 and TMPRSS2 are predominantly expressed in a transient secretory cell type in subsegmental bronchial branches. bioRxiv (2020) doi:10.1101/2020.03.13.991455.

33. Chua, R. L. et al. COVID-19 severity correlates with airway epithelium–immune cell interactions identified by single-cell analysis. Nat. Biotechnol. (2020) doi:10.1038/s41587-020-0602-4.

34. Schaefer, I. M. et al. In situ detection of SARS-CoV-2 in lungs and airways of patients with COVID-19. Mod. Pathol. (2020) doi:10.1038/s41379-020-0595-z.

35. Hou, Y. J. et al. SARS-CoV-2 Reverse Genetics Reveals a Variable Infection Gradient in the Respiratory Tract. Cell (2020) doi:10.1016/j.cell.2020.05.042.

36. Zhu, N. et al. Morphogenesis and cytopathic effect of SARS-CoV-2 infection in human airway epithelial cells. Nat. Commun. (2020) doi:10.1038/s41467-020-17796-z.

37. Ravindra, N. G. et al. Single-cell longitudinal analysis of SARS-CoV-2 infection in human airway epithelium. bioRxiv (2020) doi:10.1101/2020.05.06.081695.

38. Blanco-Melo, D. et al. Imbalanced Host Response to SARS-CoV-2 Drives Development of COVID-19. Cell (2020) doi:10.1016/j.cell.2020.04.026.

39. Cheemarla, N. R. et al. Magnitude and timing of the antiviral response determine SARS-CoV-2 replication early in infection. medRxiv (2021).

40. Mykytyn, A. Z. et al. Sars-cov-2 entry into human airway organoids is serine protease-mediated and facilitated by the multibasic cleavage site. Elife (2021) doi:10.7554/ELIFE.64508.

41. Lamers, M. M. et al. An organoid-derived bronchioalveolar model for SARS-CoV-2 infection of human alveolar type II-like cells. EMBO J. (2021) doi:10.15252/embj.2020105912.

42. Huang, J. et al. SARS-CoV-2 Infection of Pluripotent Stem Cell-Derived Human Lung Alveolar Type 2 Cells Elicits a Rapid Epithelial-Intrinsic Inflammatory Response. Cell Stem Cell (2020) doi:10.1016/j.stem.2020.09.013.

43. Pellegrini, L. et al. SARS-CoV-2 Infects the Brain Choroid Plexus and Disrupts the Blood-CSF Barrier in Human Brain Organoids. Cell Stem Cell (2020) doi:10.1016/j.stem.2020.10.001.

44. Purkayastha, A. et al. Direct Exposure to SARS-CoV-2 and Cigarette Smoke Increases Infection Severity and Alters the Stem Cell-Derived Airway Repair Response. Cell Stem Cell (2020) doi:10.1016/j.stem.2020.11.010.

45. Speranza, E. et al. Single-cell RNA sequencing reveals SARS-CoV-2 infection dynamics in lungs of African green monkeys. Sci. Transl. Med. (2021) doi:10.1126/scitranslmed.abe8146.

46. Munster, V. et al. Respiratory disease and virus shedding in rhesus macaques inoculated with SARS-CoV-2. bioRxiv (2020) doi:10.1101/2020.03.21.001628.

47. Chandrashekar, A. et al. SARS-CoV-2 infection protects against rechallenge in rhesus macaques. Science (80-. ). (2020) doi:10.1126/science.abc4776.

48. Chan, J. F. W. et al. Simulation of the Clinical and Pathological Manifestations of Coronavirus Disease 2019 (COVID-19) in a Golden Syrian Hamster Model: Implications for Disease Pathogenesis and Transmissibility. Clin. Infect. Dis. (2020) doi:10.1093/cid/ciaa325.

49. Sia, S. F. et al. Pathogenesis and transmission of SARS-CoV-2 in golden hamsters. Nature (2020) doi:10.1038/s41586-020-2342-5.

50. Sun, S. H. et al. A Mouse Model of SARS-CoV-2 Infection and Pathogenesis. Cell Host Microbe (2020) doi:10.1016/j.chom.2020.05.020.

51. Bao, L. et al. The pathogenicity of SARS-CoV-2 in hACE2 transgenic mice. Nature (2020) doi:10.1038/s41586-020-2312-y.

52. Jiang, R. Di et al. Pathogenesis of SARS-CoV-2 in Transgenic Mice Expressing Human Angiotensin-Converting Enzyme 2. Cell (2020) doi:10.1016/j.cell.2020.05.027.

53. Israelow, B. et al. Mouse model of SARS-CoV-2 reveals inflammatory role of type i interferon signaling. J. Exp. Med. (2020) doi:10.1084/JEM.20201241.

54. Kim, Y. Il et al. Infection and Rapid Transmission of SARS-CoV-2 in Ferrets. Cell Host Microbe (2020) doi:10.1016/j.chom.2020.03.023.

55. Richard, M. et al. SARS-CoV-2 is transmitted via contact and via the air between ferrets. Nat. Commun. (2020) doi:10.1038/s41467-020-17367-2.

56. Muñoz-Fontela, C. et al. Animal models for COVID-19. Nature (2020) doi:10.1038/s41586-020-2787-6.

57. Zhang, Q. et al. Inborn errors of type I IFN immunity in patients with life-threatening COVID-19. Science (80-. ). (2020) doi:10.1126/science.abd4570.

58. Bastard, P. et al. Autoantibodies against type I IFNs in patients with life-threatening COVID-19. Science (80-. ). (2020) doi:10.1126/science.abd4585.

59. Combes, A. J. et al. Global absence and targeting of protective immune states in severe COVID-19. Nature (2021) doi:10.1038/s41586-021-03234-7.

60. World Health Organization. WHO R&D Blueprint novel Coronavirus COVID-19 Therapeutic Trial Synopsis. World Heal. Organ. (2020).

61. Gierahn, T. M. et al. Seq-Well: Portable, low-cost rna sequencing of single cells at high throughput. Nat. Methods (2017) doi:10.1038/nmeth.4179.

62. Hughes, T. K. et al. Highly Efficient, Massively-Parallel Single-Cell RNA-Seq Reveals Cellular States and Molecular Features of Human Skin Pathology. bioRxiv (2019) doi:10.1101/689273.

63. Ordovas-Montanes, J. et al. Allergic inflammatory memory in human respiratory epithelial progenitor cells. Nature (2018) doi:10.1038/s41586-018-0449-8.

64. Garcıá, S. R. et al. Novel dynamics of human mucociliary differentiation revealed by single-cell RNA sequencing of nasal epithelial cultures. Dev. (2019) doi:10.1242/dev.177428.

65. Deprez, M. et al. A single-cell atlas of the human healthy airways. Am. J. Respir. Crit. Care Med. (2020) doi:10.1164/rccm.201911-2199OC.

66. Montoro, D. T. et al. A revised airway epithelial hierarchy includes CFTR-expressing ionocytes. Nature (2018) doi:10.1038/s41586-018-0393-7.

67. Plasschaert, L. W. et al. A single-cell atlas of the airway epithelium reveals the CFTR-rich pulmonary ionocyte. Nature (2018) doi:10.1038/s41586-018-0394-6.

68. Basak, O. et al. Induced Quiescence of Lgr5+ Stem Cells in Intestinal Organoids Enables Differentiation of Hormone-Producing Enteroendocrine Cells. Cell Stem Cell (2017) doi:10.1016/j.stem.2016.11.001.

69. Hoffmann, M. et al. SARS-CoV-2 Cell Entry Depends on ACE2 and TMPRSS2 and Is Blocked by a Clinically Proven Protease Inhibitor. Cell (2020) doi:10.1016/j.cell.2020.02.052.

70. Li, W. et al. Angiotensin-converting enzyme 2 is a functional receptor for the SARS coronavirus. Nature (2003) doi:10.1038/nature02145.

71. Yan, R. et al. Structural basis for the recognition of SARS-CoV-2 by full-length human ACE2. Science (80-. ). (2020) doi:10.1126/science.abb2762.

72. Wrapp, D. et al. Cryo-EM structure of the 2019-nCoV spike in the prefusion conformation. Science (80-. ). (2020) doi:10.1126/science.aax0902.

73. Wang, Q. et al. Structural and Functional Basis of SARS-CoV-2 Entry by Using Human ACE2. Cell (2020) doi:10.1016/j.cell.2020.03.045.

74. Raj, V. S. et al. Dipeptidyl peptidase 4 is a functional receptor for the emerging human coronavirus-EMC. Nature (2013) doi:10.1038/nature12005.

75. Yeager, C. L. et al. Human aminopeptidase N is a receptor for human coronavirus 229E. Nature (1992) doi:10.1038/357420a0.

76. Bochkov, Y. A. et al. Cadherin-related family member 3, a childhood asthma susceptibility gene product, mediates rhinovirus C binding and replication. Proc. Natl. Acad. Sci. U. S. A. (2015) doi:10.1073/pnas.1421178112.

77. Bergen, V., Lange, M., Peidli, S., Wolf, F. A. & Theis, F. J. Generalizing RNA velocity to transient cell states through dynamical modeling. Nat. Biotechnol. (2020) doi:10.1038/s41587-020-0591-3.

78. La Manno, G. et al. RNA velocity of single cells. Nature (2018) doi:10.1038/s41586-018-0414-6.

79. Tata, P. R. et al. Dedifferentiation of committed epithelial cells into stem cells in vivo. Nature (2013) doi:10.1038/nature12777.

80. Cosgrove, P. R., Redhu, N. S., Tang, Y., Monuteaux, M. C. & Horwitz, B. H. Characterizing T cell subsets in the nasal mucosa of children with acute respiratory symptoms. Pediatr. Res. (2021) doi:10.1038/s41390-021-01364-2.

81. Blume, C. et al. A novel ACE2 isoform is expressed in human respiratory epithelia and is upregulated in response to interferons and RNA respiratory virus infection. Nat. Genet. (2021) doi:10.1038/s41588-020-00759-x.

82. Ng, K. W. et al. Tissue-specific and interferon-inducible expression of nonfunctional ACE2 through endogenous retroelement co-option. Nat. Genet. (2020) doi:10.1038/s41588-020-00732-8.

83. Onabajo, O. O. et al. Interferons and viruses induce a novel truncated ACE2 isoform and not the full-length SARS-CoV-2 receptor. Nat. Genet. (2020) doi:10.1038/s41588-020-00731-9.

84. Cao, Y. et al. Single-cell analysis of upper airway cells reveals host-viral dynamics in influenza infected adults. bioRxiv (2020) doi:10.1101/2020.04.15.042978.

85. Lemieux, J. E. et al. Phylogenetic analysis of SARS-CoV-2 in Boston highlights the impact of superspreading events. Science (80-. ). (2020) doi:10.1126/science.abe3261.

86. Wood, D. E., Lu, J. & Langmead, B. Improved metagenomic analysis with Kraken 2. Genome Biol. (2019) doi:10.1186/s13059-019-1891-0.

87. Kim, D. et al. The Architecture of SARS-CoV-2 Transcriptome. Cell (2020) doi:10.1016/j.cell.2020.04.011.

88. Delorey, T. et al. A single-cell and spatial atlas of autopsy tissues reveals pathology and cellular targets of SARS-CoV-2. bioRxiv (2021).

89. Garcia-Beltran, W. F. et al. COVID-19-neutralizing antibodies predict disease severity and survival. Cell (2021) doi:10.1016/j.cell.2020.12.015.

90. Zohar, T. et al. Compromised Humoral Functional Evolution Tracks with SARS-CoV-2 Mortality. Cell (2020) doi:10.1016/j.cell.2020.10.052.

91. Long, Q. X. et al. Antibody responses to SARS-CoV-2 in patients with COVID-19. Nat. Med. (2020) doi:10.1038/s41591-020-0897-1.

92. Kotliar, D. et al. Single-Cell Profiling of Ebola Virus Disease In Vivo Reveals Viral and Host Dynamics. Cell (2020) doi:10.1016/j.cell.2020.10.002.

93. Fleming, S. J., Marioni, J. C. & Babadi, M. CellBender remove-background: A deep generative model for unsupervised removal of background noise from scRNA-seq datasets. bioRxiv (2019) doi:10.1101/791699.

94. Hu, B., Guo, H., Zhou, P. & Shi, Z. L. Characteristics of SARS-CoV-2 and COVID-19. Nature Reviews Microbiology (2020) doi:10.1038/s41579-020-00459-7.

95. Fung, T. S. & Liu, D. X. Human Coronavirus: Host-Pathogen Interaction. Annu. Rev. Microbiol. (2019) doi:10.1146/annurev-micro-020518-115759.

96. Sawicki, S. G., Sawicki, D. L. & Siddell, S. G. A Contemporary View of Coronavirus Transcription. J. Virol. (2007) doi:10.1128/jvi.01358-06.

97. Krähling, V., Stein, D. A., Spiegel, M., Weber, F. & Mühlberger, E. Severe Acute Respiratory Syndrome Coronavirus Triggers Apoptosis via Protein Kinase R but Is Resistant to Its Antiviral Activity. J. Virol. (2009) doi:10.1128/jvi.01245-08.

98. Zhao, X. et al. Interferon induction of IFITM proteins promotes infection by human coronavirus OC43. Proc. Natl. Acad. Sci. U. S. A. (2014) doi:10.1073/pnas.1320856111.

99. Daniloski, Z. et al. Identification of Required Host Factors for SARS-CoV-2 Infection in Human Cells. Cell (2021) doi:10.1016/j.cell.2020.10.030.

100. Wang, R. et al. Genetic Screens Identify Host Factors for SARS-CoV-2 and Common Cold Coronaviruses. Cell (2021) doi:10.1016/j.cell.2020.12.004.

101. Wei, J. et al. Genome-wide CRISPR Screens Reveal Host Factors Critical for SARS-CoV-2 Infection. Cell (2021) doi:10.1016/j.cell.2020.10.028.

102. Schneider, W. M. et al. Genome-Scale Identification of SARS-CoV-2 and Pan-coronavirus Host Factor Networks. Cell (2021) doi:10.1016/j.cell.2020.12.006.

103. Ordovas-Montanes, J., Beyaz, S., Rakoff-Nahoum, S. & Shalek, A. K. Distribution and storage of inflammatory memory in barrier tissues. Nat. Rev. Immunol. (2020) doi:10.1038/s41577-019-0263-z.

104. Kamitani, W., Huang, C., Narayanan, K., Lokugamage, K. G. & Makino, S. A two-pronged strategy to suppress host protein synthesis by SARS coronavirus Nsp1 protein. Nat. Struct. Mol. Biol. (2009) doi:10.1038/nsmb.1680.

105. Lokugamage, K. G. et al. Middle East Respiratory Syndrome Coronavirus nsp1 Inhibits Host Gene Expression by Selectively Targeting mRNAs Transcribed in the Nucleus while Sparing mRNAs of Cytoplasmic Origin. J. Virol. (2015) doi:10.1128/jvi.01352-15.

106. Knoops, K. et al. SARS-coronavirus replication is supported by a reticulovesicular network of modified endoplasmic reticulum. PLoS Biol. (2008) doi:10.1371/journal.pbio.0060226.

107. Menachery, V. D. et al. Pathogenic influenza viruses and coronaviruses utilize similar and contrasting approaches to control interferon-stimulated gene responses. MBio (2014) doi:10.1128/mBio.01174-14.

108. Konno, Y. et al. SARS-CoV-2 ORF3b Is a Potent Interferon Antagonist Whose Activity Is Increased by a Naturally Occurring Elongation Variant. Cell Rep. (2020) doi:10.1016/j.celrep.2020.108185.

109. Banerjee, A. K. et al. SARS-CoV-2 Disrupts Splicing, Translation, and Protein Trafficking to Suppress Host Defenses. Cell (2020) doi:10.1016/j.cell.2020.10.004.

110. Snijder, E. J. et al. A unifying structural and functional model of the coronavirus replication organelle: Tracking down RNA synthesis. PLoS Biol. (2020) doi:10.1371/journal.pbio.3000715.

111. Broggi, A., Granucci, F. & Zanoni, I. Type III interferons: Balancing tissue tolerance and resistance to pathogen invasion. J. Exp. Med. (2020) doi:10.1084/jem.20190295.

112. Major, J. et al. Type I and III interferons disrupt lung epithelial repair during recovery from viral infection. Science (80-. ). (2020) doi:10.1126/science.abc2061.

113. Silvin, A. et al. Elevated Calprotectin and Abnormal Myeloid Cell Subsets Discriminate Severe from Mild COVID-19. Cell (2020) doi:10.1016/j.cell.2020.08.002.

114. Bruchez, A. et al. MHC class II transactivator CIITA induces cell resistance to ebola virus and SARS-like coronaviruses. Science (80-. ). (2020) doi:10.1126/science.abb3753.

115. Feld, J. J. et al. Peginterferon-lambda for the treatment of COVID-19 in outpatients. medRxiv (2020) doi:10.1101/2020.11.09.20228098.

116. Monk, P. D. et al. Safety and efficacy of inhaled nebulised interferon beta-1a (SNG001) for treatment of SARS-CoV-2 infection: a randomised, double-blind, placebo-controlled, phase 2 trial. Lancet Respir. Med. (2020) doi:10.1016/S2213-2600(20)30511-7.

117. Wang, N. et al. Retrospective Multicenter Cohort Study Shows Early Interferon Therapy Is Associated with Favorable Clinical Responses in COVID-19 Patients. Cell Host Microbe (2020) doi:10.1016/j.chom.2020.07.005.

118. Hoagland, D. A. et al. Leveraging the antiviral type-I interferon system as a first line defense against SARS-CoV-2 pathogenicity. Immunity (2021) doi:10.1016/j.immuni.2021.01.017.

119. Tang, Y. et al. Human Nasopharyngeal Swab Processing for Viable Single-Cell Suspension. protocols.io (2020).

120. Aicher, T. P. et al. Seq-Well: A sample-efficient, portable picowell platform for massively parallel single-cell RNA sequencing. in Methods in Molecular Biology (2019). doi:10.1007/978-1-4939-9240-9_8.

121. Macosko, E. Z. et al. Highly parallel genome-wide expression profiling of individual cells using nanoliter droplets. Cell (2015) doi:10.1016/j.cell.2015.05.002.

122. Li, B. et al. Cumulus provides cloud-based data analysis for large-scale single-cell and single-nucleus RNA-seq. Nat. Methods (2020) doi:10.1038/s41592-020-0905-x.

123. Petukhov, V. et al. dropEst: Pipeline for accurate estimation of molecular counts in droplet-based single-cell RNA-seq experiments. Genome Biol. (2018) doi:10.1186/s13059-018-1449-6.

124. Stuart, T. et al. Comprehensive Integration of Single-Cell Data. Cell (2019) doi:10.1016/j.cell.2019.05.031.

125. McDavid, A. et al. Data exploration, quality control and testing in single-cell qPCR-based gene expression experiments. Bioinformatics (2013) doi:10.1093/bioinformatics/bts714.

126. Love, M. I., Anders, S. & Huber, W. Differential analysis of count data - the DESeq2 package. Genome Biology (2014).

127. Huang, D. W., Sherman, B. T. & Lempicki, R. A. Systematic and integrative analysis of large gene lists using DAVID bioinformatics resources. Nat. Protoc. (2009) doi:10.1038/nprot.2008.211.

128. Korotkevich, G. et al. Fast gene set enrichment analysis. bioRxiv (2021) doi:10.1101/060012.

129. R Core Team. R: A language and environment for statistical computing. R Foundation for Statistical Computing (2019).

130. Wickham, H. ggplot2: Elegant Graphics for Data Analysis. Journal of the Royal Statistical Society: Series A (Statistics in Society) (2016).

131. Butler, A., Hoffman, P., Smibert, P., Papalexi, E. & Satija, R. Integrating single-cell transcriptomic data across different conditions, technologies, and species. Nat. Biotechnol. (2018) doi:10.1038/nbt.4096.

132. Gu, Z., Eils, R. & Schlesner, M. Complex heatmaps reveal patterns and correlations in multidimensional genomic data. Bioinformatics (2016) doi:10.1093/bioinformatics/btw313.

133. Gu, Z., Gu, L., Eils, R., Schlesner, M. & Brors, B. Circlize implements and enhances circular visualization in R. Bioinformatics (2014) doi:10.1093/bioinformatics/btu393.

134. Ballestar, E. et al. Single cell profiling of COVID-19 patients: An international data resource from multiple tissues. medRxiv (2020) doi:10.1101/2020.11.20.20227355.

