## Supplementary Figures for "Impaired local intrinsic immunity to SARS-CoV-2 infection in severe COVID-19"

#### Supplementary Material

Carly G. K. Ziegler[1,2,3,4,5,*], Vincent N. Miao[1,2,3,5,*], Anna H. Owings[6,*], Andrew W. Navia[2,3,5,7,*], Ying Tang[8,*], Joshua D. Bromley[2,3,5,9,*], Peter Lotfy[3,8], Meredith Sloan[6], Hannah Laird[10], Haley B. Williams[10], Micayla George[2,3,5], Riley S. Drake[2,3,5], Taylor Christian[6], Adam Parker[10], Campbell B. Sindel[11], Molly W. Burger[12], Yilianys Pride[10], Mohammad Hasan[10], George E. Abraham III[11], Michal Senitko[11], Tanya O. Robinson[10], Alex K. Shalek[1,2,3,4,5,7,13,15,16,^], Sarah C. Glover[10,^], Bruce H. Horwitz[8,14,15,^], Jose Ordovas-Montanes[3,8,15,16,^]

[1] Program in Health Sciences & Technology, Harvard Medical School & MIT, Boston, MA 02115, USA
[2] Ragon Institute of MGH, MIT, and Harvard, Cambridge, MA 02139, USA
[3] Broad Institute of MIT and Harvard, Cambridge, MA 02142, USA
[4] Harvard Graduate Program in Biophysics, Harvard University, Cambridge, MA 02138, USA
[5] Institute for Medical Engineering & Science, Massachusetts Institute of Technology, Cambridge, MA 02139, USA
[6] Department of Medicine, University of Mississippi Medical Center, Jackson, MS 39216, USA
[7] Department of Chemistry, Massachusetts Institute of Technology, Cambridge, MA 02139, USA
[8] Division of Gastroenterology, Hepatology, and Nutrition, Boston Children's Hospital, Boston, MA 02115, USA
[9] Department of Microbiology, Massachusetts Institute of Technology, Cambridge, MA 02139, USA
[10] Division of Digestive Diseases, University of Mississippi Medical Center, Jackson, MS 39216, USA
[11] Division of Pulmonary, Critical Care, and Sleep Medicine, University of Mississippi Medical Center, Jackson, MS 39216, USA
[12] Obstetrics and Gynecology, University of Mississippi Medical Center, Jackson, MS 39216, USA
[13] Koch Institute for Integrative Cancer Research, Massachusetts Institute of Technology, Cambridge, MA 02139, USA
[14] Division of Emergency Medicine, Boston Children's Hospital, Boston, MA 02115, USA
[15] Program in Immunology, Harvard Medical School, Boston, MA 02115, USA
[16] Harvard Stem Cell Institute, Cambridge, MA 02138, USA
* these authors contributed equally
^ these senior authors contributed equally

Correspondence to: Jose Ordovas-Montanes, Bruce Horwitz, Sarah C. Glover, and Alex K. Shalek

### Supplementary Figure 1

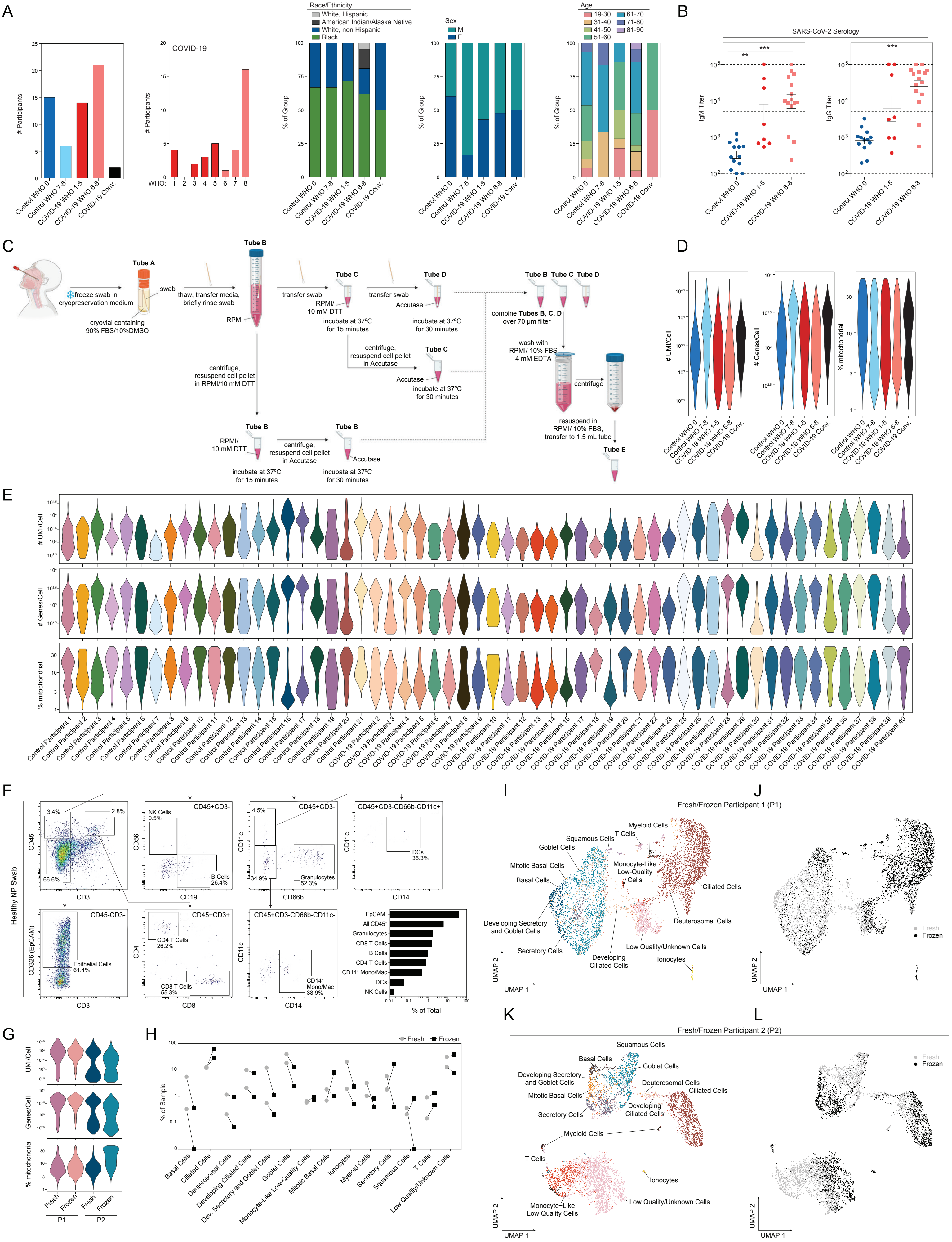

**Supplementary Figure 1. Cohort and Cellular Composition of Nasopharyngeal Swabs**

*Related to Figure 1, Table 1*

- **A.** Cohort composition and participant demographics (see also **Table 1**).
- **B.** SARS-CoV-2 serology: IgM (left) and IgG (right) titers from a subset of Control WHO 0 (blue circles, n=13) and COVID-19 (red circles, mild/moderate: n=8; pink squares, severe: n=15) participants. Plasma samples taken on same day of nasopharyngeal swab. Statistical testing by Kruskal-Wallis test with Dunn's post hoc testing. Asterisks represent results from Dunn's test: ** $p < 0.01$, *** $p < 0.001$. Dashed lines: lower limit of detection: 100; upper limit of detection: 100,000; positive threshold: 5,000.
- **C.** Detailed schematic of sample preparation and cell processing from nasal swabs (created with BioRender).
- **D.** Single-cell quality metrics by group (after filtering for low-quality cells, see **Methods**).
- **E.** Single-cell quality metrics by participant (after filtering for low quality cells).
- **F.** Flow cytometry and gating scheme of cells from a representative fresh nasopharyngeal swab from a healthy participant. Bottom right: quantification of cellular proportions.
- **G.** Quality metrics for matched fresh vs. frozen nasal swabs from two healthy participants (P1 and P2).
- **H.** Percent composition of each cell type by processing type: fresh (grey circles) or frozen (black squares).
- **I.** UMAP of cell types from P1.
- **J.** UMAP from P1 as in **I**, colored by fresh (grey) vs. frozen (black).
- **K.** UMAP of cell types from P2.
- **L.** UMAP from P2 as in **K**, colored by fresh (grey) vs. frozen (black).

### Supplementary Figure 2

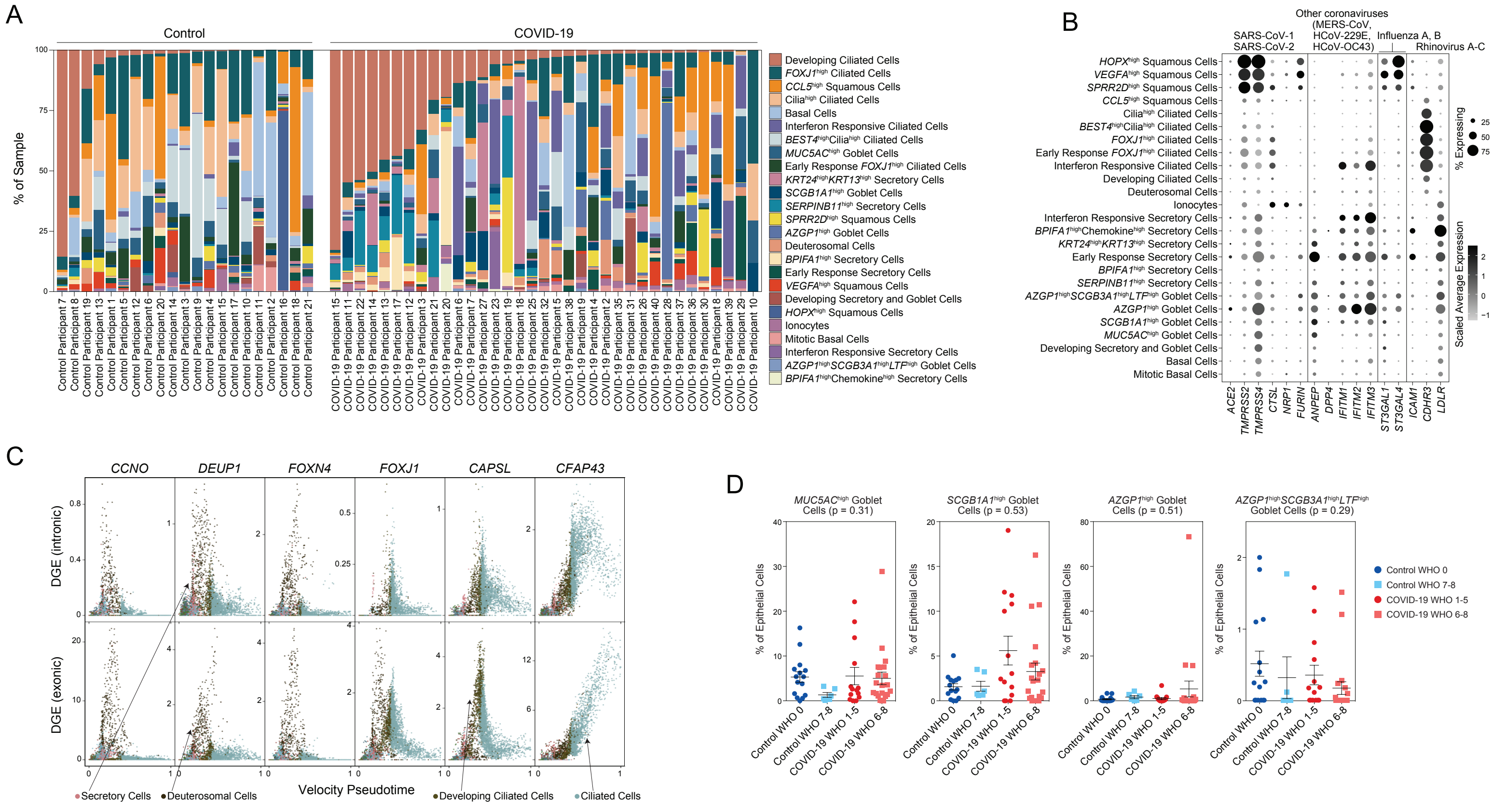

**Supplementary Figure 2. Epithelial Diversity and Differentiation in the Nasopharyngeal Mucosa During COVID-19**

*Related to Figure 2*

- **A.** Proportional abundance of detailed epithelial cell types by participant. Ordered within group by developing ciliated cell proportion.
- **B.** Expression of entry factors for SARS-CoV-2 and other common upper respiratory viruses among detailed epithelial cell types. Dot size represents fraction of cell type (rows) expressing a given gene (columns). Dot hue represents scaled average expression by gene column.
- **C.** Plot of gene expression by epithelial cell velocity pseudotime (over all epithelial cells). Select genes significantly associated with ciliated cell pseudotime (FDR < 0.01). Points colored by coarse cell type annotations. Top: alignment to unspliced (intronic) regions. Bottom: alignment to spliced (exonic) regions.
- **D.** Proportion of goblet cell subtypes (detailed annotation) by sample, normalized to all epithelial cells. Statistical test above graph represents Kruskal-Wallis test results across all groups (following FDR correction).

### Supplementary Figure 3

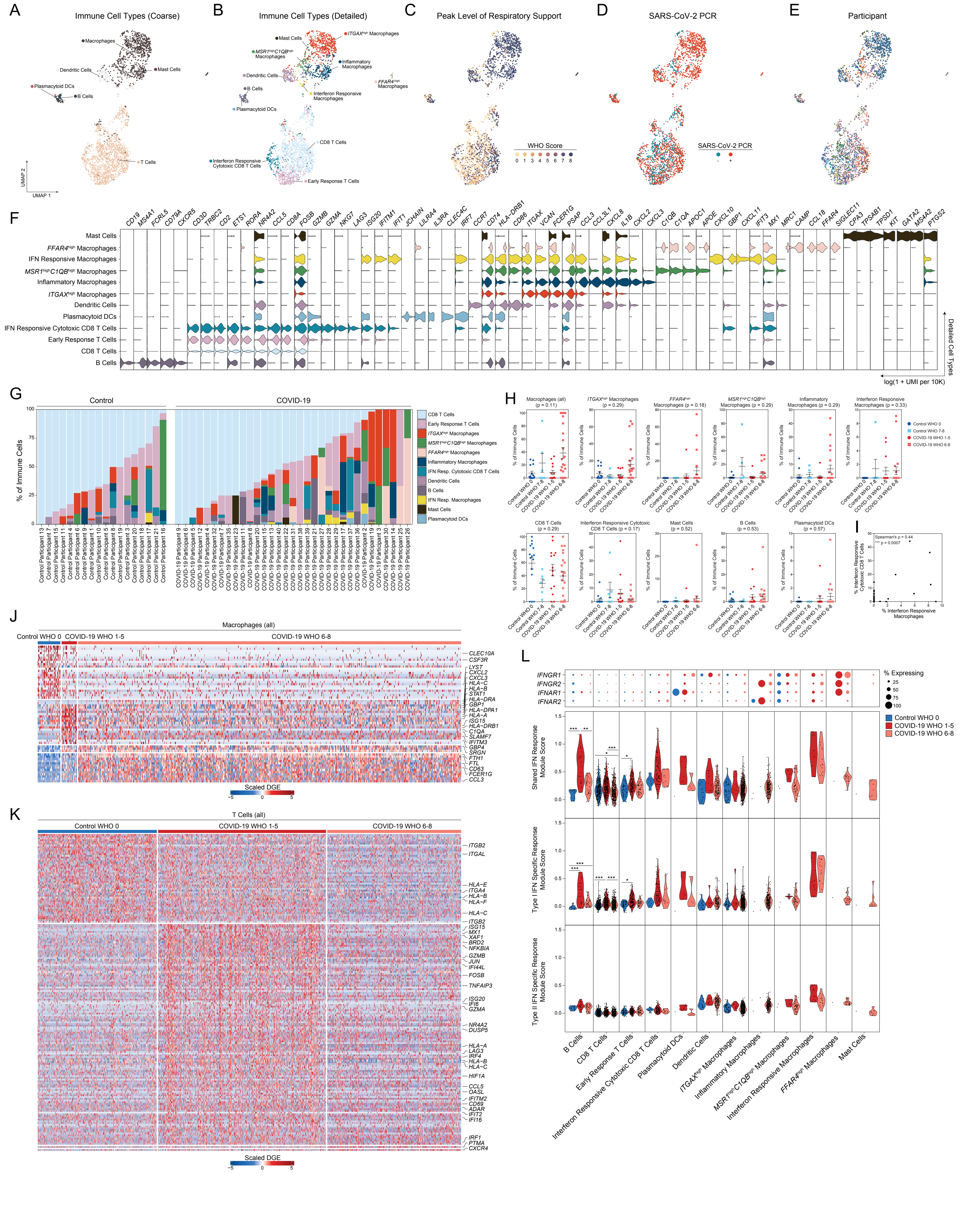

**Supplementary Figure 3. Immune Cell Diversity in the Nasopharyngeal Mucosa During COVID-19**

*Related to Figures 1 and 3*

- **A.** UMAP of 3,640 immune cells following re-clustering, colored by coarse cell types.
- **B.** UMAP as in **A**, colored by detailed cell annotations.
- **C.** UMAP as in **A**, colored by peak level of respiratory support (WHO illness severity scale).
- **D.** UMAP as in **A**, colored by SARS-CoV-2 diagnostic PCR status.
- **E.** UMAP as in **A**, colored by participant.
- **F.** Violin plots (log(1+normalized UMI per 10k)) of cluster marker genes (FDR < 0.01) for detailed immune cell type annotations (as in **B**).
- **G.** Proportional abundance of detailed immune cell types by participant. Ordered within group by CD8 T cell proportion.
- **H.** Proportion of immune cell subtypes by sample and disease group, normalized to all immune cells. Statistical test above graph represents Kruskal-Wallis test results across all cell types (following FDR correction).
- **I.** Proportion of interferon responsive macrophages vs. proportion of interferon responsive cytotoxic CD8 T cells per sample, normalized to total immune cells. Including all samples, Control and COVID-19 groups.
- **J.** Heatmap of significantly DE genes between macrophages (all, coarse annotation) from different disease groups. Values represent row(gene)-scaled digital gene expression (DGE) following log(1+UMI per 10K) normalization.
- **K.** Heatmap of significantly DE genes between T cells (all, coarse annotation) from different disease groups. Values represent row(gene)-scaled digital gene expression (DGE) following log(1+UMI per 10K) normalization.
- **L.** Top: Dot plot of *IFNGR1, IFNGR2*, *IFNAR1,* and *IFNAR2* gene expression among all detailed immune subtypes. Bottom: Violin plots of module scores, split by Control WHO 0 (blue), COVID-19 WHO 1-5 (red), and COVID-19 WHO 6-8 (pink). Gene modules represent transcriptional responses of human basal cells from the nasal epithelium following *in vitro* treatment with IFN$\alpha$ or IFN$\gamma$. Significance by Wilcoxon signed-rank test. P-values following Bonferroni-correction: * p< 0.05, ** p < 0.01, *** p < 0.001.

#### Supplementary Figure 4

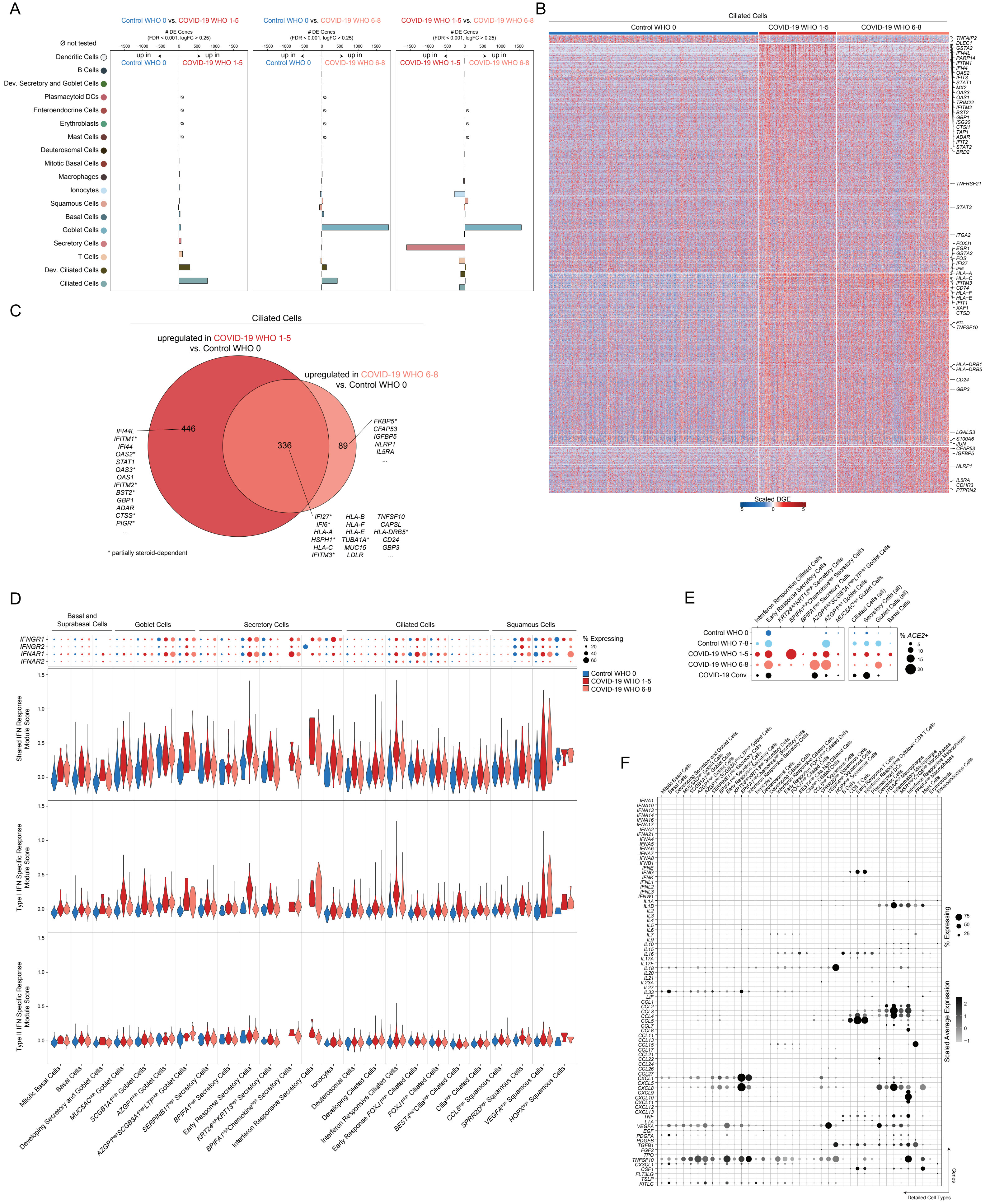

**Supplementary Figure 4. Cell-Type Specific and Shared Transcriptional Responses to SARS-CoV-2 Infection**

*Related to Figure 3*

- **A.** Abundance of significant DE genes by coarse cell type between Control WHO 0 and COVID-19 WHO 1-5 samples (left), Control WHO 0 and COVID-19 WHO 6-8 samples (middle) and COVID-19 WHO 1-5 vs. COVID-19 WHO 6-8 samples (right). Gene significance cutoffs: FDR-corrected $p < 0.001$, log2 fold change > 0.25.
- **B.** Heatmap of significantly DE genes between ciliated cells (all, coarse annotation) from different disease groups. Values represent row(gene)-scaled digital gene expression (DGE) following log(1+UMI per 10K) normalization.
- **C.** Venn diagram of significantly upregulated genes among ciliated cells between COVID-19 WHO 1-5 vs. Control WHO 0 (red) and COVID-19 WHO 6-8 vs. Control WHO 0 (pink). Asterisk: genes impacted by corticosteroid treatment within each group.
- **D.** Top: Dot plot of *IFNGR1, IFNGR2*, *IFNAR1,* and *IFNAR2* gene expression among all detailed epithelial subtypes. Bottom: Violin plots of module scores, split by Control WHO 0 (blue), COVID-19 WHO 1-5 (red), and COVID-19 WHO 6-8 (pink). Gene modules represent transcriptional responses of human basal cells from the nasal epithelium following *in vitro* treatment with IFN$\alpha$ or IFN$\gamma$. Significance by Wilcoxon signed-rank test. P-values following Bonferroni-correction: * $p< 0.05$, ** $p < 0.01$, *** $p < 0.001$.
- **E.** Dot plot of *ACE2* expression across select epithelial cell types and subsets.
- **F.** Dot plot of interferon and cytokine expression among detailed epithelial and immune cell types.

### Supplementary Figure 5

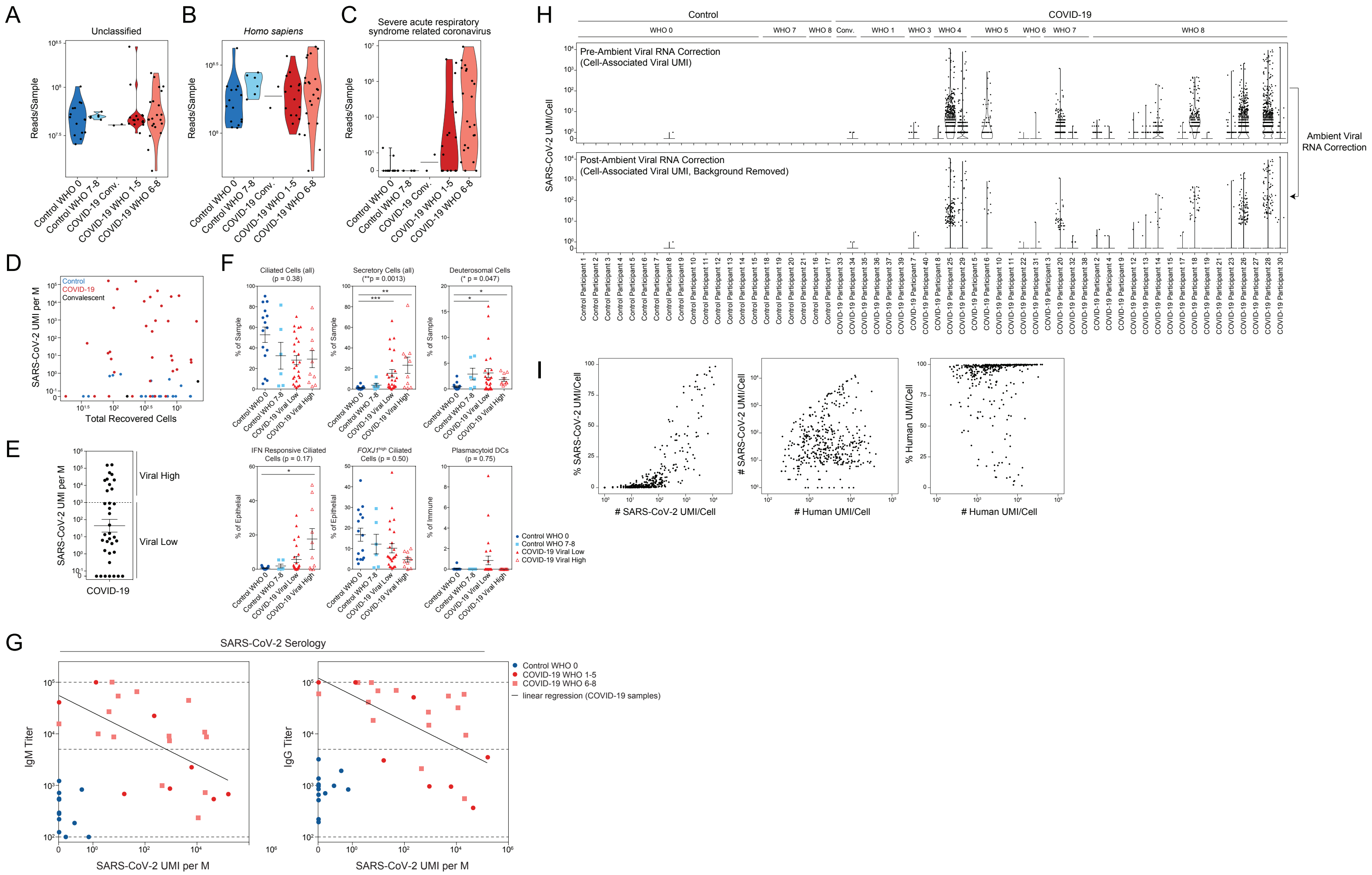

**Supplementary Figure 5. Detection of SARS-CoV-2 RNA from Single-Cell RNA-Seq Data**

*Related to Figures 4 and 5*

- **A.** Metatranscriptomic classification of all scRNA-seq reads using Kraken2: reads per sample annotated as unclassified.
- **B.** Metatranscriptomic classification of all scRNA-seq reads using Kraken2: reads per sample annotated as *Homo sapiens.*
- **C.** Metatranscriptomic classification of all scRNA-seq reads using Kraken2: reads per sample annotated as SARS-related coronaviruses.
- **D.** Total recovered cells per sample vs. normalized abundance of SARS-CoV-2 aligning UMI from all scRNA-seq UMI (including those derived from ambient/low-complexity cell barcodes).
- **E.** Normalized abundance of SARS-CoV-2 aligning UMI across all COVID-19 participants. Dashed line represents partition between “Viral High” vs “Viral Low” samples (1,000 SARS-CoV-2 UMI/million (M) UMI).
- **F.** Proportional abundance of selected cell types according to total SARS-CoV-2 abundance among COVID-19 samples, stratified by cutoffs in panel **E**. Statistical test above graph represents FDR-corrected Kruskal-Wallis test statistic across all groups. Statistical significance asterisks within box represent significant results from Dunn’s post-hoc testing. * $p < 0.05$, ** $p < 0.01$, *** $p < 0.001$.
- **G.** Normalized abundance of SARS-CoV-2 aligning UMI vs. anti-SARS-CoV-2 IgM (left) or IgG titers (right). Plasma samples taken on same day of nasopharyngeal swab. Subset of Control WHO 0 (blue circles, n=13) and COVID-19 (red circles, mild/moderate: n=8; pink squares, severe: n=15) participants. Dashed lines: lower limit of detection: 100; upper limit of detection: 100,000; positive threshold: 5,000. Pearson’s correlation of COVID-19 samples: IgM: $r = -0.59$, ** $p = 0.0028$; IgG: $r = -0.60$, ** $p = 0.0025$.
- **H.** Abundance of SARS-CoV-2 aligning UMI/cell by participant prior to (top) and following (bottom) ambient viral RNA correction (see **Methods**).
- **I.** Quality metrics among 415 SARS-CoV-2 RNA+ cells (associated with high-quality cell barcodes and following ambient viral RNA correction). Left: abundance of SARS-CoV-2 aligning UMI vs. percent of all SARS-CoV-2 aligned reads (per cell barcode). Middle: abundance of human (GRCh38)-aligning UMI vs. abundance of SARS-CoV-2 aligning UMI. Right: abundance of human (GRCh38) aligning UMI vs. percent of all human aligned reads (per cell barcode).

### Supplementary Figure 6

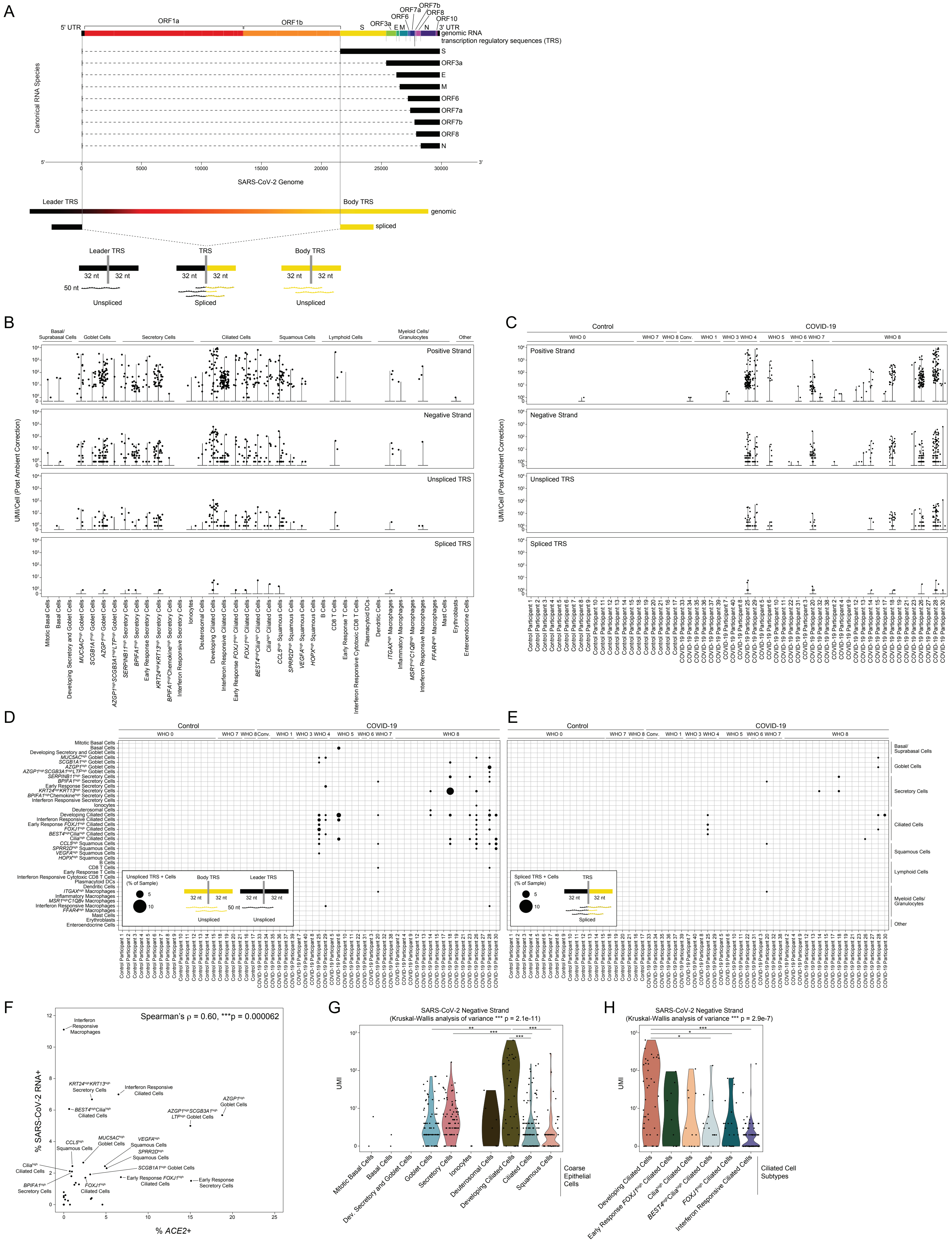

**Supplementary Figure 6. SARS-CoV-2 RNA and Cell Types Containing Viral Reads**

*Related to Figures 4 and 5*

- **A.** Schematic of method to distinguish unspliced from spliced SARS-CoV-2 RNA species by searching for reads which align across a spliced or genomic Transcription Regulatory Sequence (TRS).
- **B.** Abundance of SARS-CoV-2 aligning UMI/Cell per detailed cell type (following ambient viral RNA correction), split by UMI aligning to the viral positive strand, negative strand, 70-mer region across an unspliced TRS, and 70-mer region across a spliced TRS.
- **C.** Abundance of SARS-CoV-2 aligning UMI/Cell per participant (following ambient viral RNA correction), split by UMI aligning to the viral positive strand, negative strand, 70-mer region across an unspliced TRS, and 70-mer region across a spliced TRS.
- **D.** Dot plot of SARS-CoV-2 unspliced TRS aligning UMI by participant (columns) and detailed cell type (rows). Dot size corresponds to the percent of cells within each sample/cell type containing unspliced TRS UMI.
- **E.** Dot plot of SARS-CoV-2 spliced TRS aligning UMI by participant (columns) and detailed cell type (rows). Dot size corresponds to the percent of cells within each sample/cell type containing spliced TRS UMI.
- **F.** Percent *ACE2*+ cells vs. percent SARS-CoV-2 RNA+ cells (after ambient correction) by detailed cell type. Including only cells from COVID-19 participants. Statistical testing using Spearman's correlation.
- **G.** Abundance of SARS-CoV-2 negative strand aligning reads by coarse epithelial cell types. Statistical significance by Kruskal-Wallis test (p-value outside box). Asterisks within box: pairwise Wilcoxon rank sum test, Bonferroni-corrected: *** $p < 0.001$, ** $p < 0.01$, * $p < 0.05$
- **H.** Abundance of SARS-CoV-2 negative strand aligning reads by detailed ciliated cell subtypes. Statistical significance by Kruskal-Wallis test (p-value outside box). Asterisks within box: pairwise Wilcoxon rank sum test, Bonferroni-corrected: *** $p < 0.001$, ** $p < 0.01$, * $p < 0.05$

Supplementary Figure 7

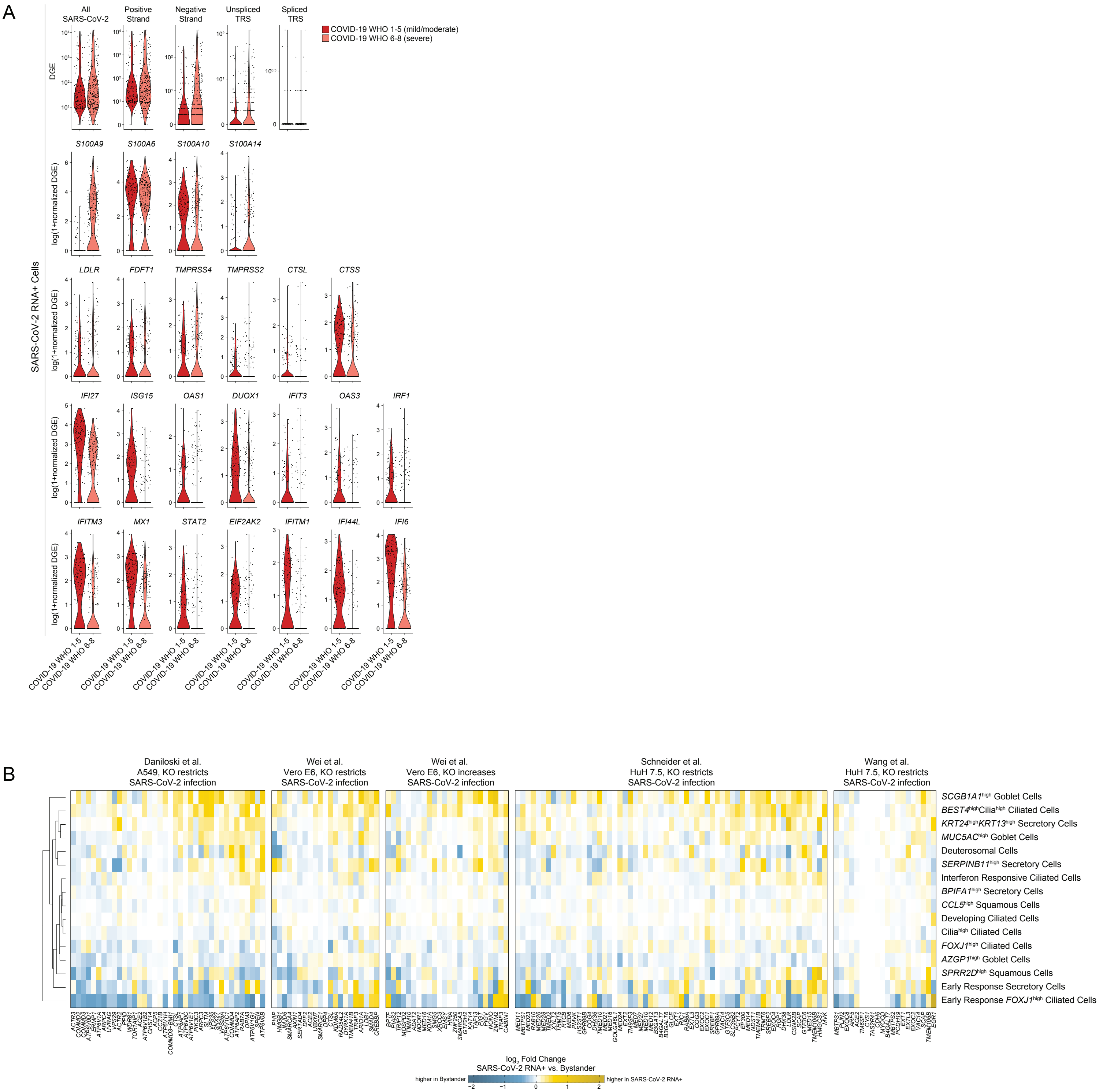

**Supplementary Figure 7. Intrinsic and Bystander Responses to SARS-CoV-2 Infection**

*Related to Figure 6*

- **A.** Violin plots of select genes upregulated in SARS-CoV-2 RNA+ cells when compared to matched bystanders. Plotting only SARS-CoV-2 RNA+ cells from COVID-19 WHO 1-5 participants (red) and COVID-19 WHO 6-8 participants (pink). Top row: SARS-CoV-2 RNA expression by alignment type.
- **B.** Heatmaps of log fold changes between SARS-CoV-2 RNA+ cells and bystander cells by cell type. Gene sets derived from four CRISPR screens for important host factors in the SARS-CoV-2 viral life cycle. Restricted to cell types with at least 5 SARS-CoV-2 RNA+ cells. Yellow: upregulated among SARS-CoV-2 RNA+ cells, blue: upregulated among bystander cells.

#### SUPPLEMENTARY TABLE LEGENDS

**Supplementary Table 1. Cell Type Marker Genes**
*Related to Figures 1, 2, Supplementary Figure 3*

**Supplementary Table 2. Differentially Expressed Genes Between Cell Types from Control WHO 0 vs. COVID-19 WHO 1-5 (Mild/Moderate)**
*Related to Figure 3*

**Supplementary Table 3. Differentially Expressed Genes Between Cell Types from Control WHO 0 vs. COVID-19 WHO 6-8 (Severe)**
*Related to Figure 3*

**Supplementary Table 4. Differentially Expressed Genes Between Cell Types from COVID-19 WHO 1-5 (Mild/Moderate) vs. COVID-19 WHO 6-8 (Severe)**
*Related to Figure 3*

**Supplementary Table 5. Common Differentially Expressed Genes Between SARS-CoV-2 RNA+ Cells and Bystander Cells**
*Related to Figure 6*
